# The effects of supergene evolution on the structure and stability of the G-matrix

**DOI:** 10.64898/2026.09.07.749873

**Authors:** Vitor Sudbrack, Charles Mullon

## Abstract

The additive genetic variances and covariances of traits, collected in the **G**-matrix, summarise heritable variation within populations and are commonly used to predict the rate and direction of short-term multivariate evolution. These (co)variances are shaped by pleiotropy and genetic linkage, two features often associated with supergenes formed by chromosomal inversions. Yet how supergene evolution affects the structure and temporal stability of the **G**-matrix remains unclear. Here, we use mathematical analysis and individual-based simulations to investigate the evolution and genetic consequences of inversions capturing multiple pleiotropic loci underlying two traits subject to disruptive and correlational selection (selection favouring particular combinations of trait values). We show that inversions evolve under disruptive selection and are maintained by balancing selection because they preserve associations among alleles that together generate discrete phenotypic morphs. The genetic architecture reflects how selection acts on the traits: selection favouring diversification along a joint trait combination generally produces a single multi-trait supergene, whereas selection favouring independent diversification of each trait produces multiple independently segregating inversions. By suppressing recombination, these inversions increase additive genetic variance and narrow-sense heritability, and stabilise the overall amount of additive genetic variation through time. When dominance is allowed to evolve, dominance relationships become coordinated across linked loci within supergenes, although most genetic variance remains additive at the population level. Together, these results link the form of multivariate selection to the evolution of supergenes and to their consequences for the structure and temporal stability of the **G**-matrix.

## 1 Introduction

The **G**-matrix collects the additive genetic variances and covariances of multiple traits (Lande, 1979, 1984; see Box 1 for the decomposition of phenotypic variance into additive, non-additive genetic and environmental components). Its diagonal elements give additive genetic variance in each trait, whereas its off-diagonal elements give additive genetic covariance between traits. These covariances can arise from pleiotropy, linkage disequilibrium, or shared developmental and physiological processes that couple traits genetically. Genetic covariances determine whether selection on one trait produces correlated responses in others (Lande, 1979; Lande and Arnold, 1983). Specifically, the short-term evolutionary response of the vector of trait means is given by the product of the **G**-matrix and the vector of selection gradients (Lande, 1979; p. 500 in Walsh and Lynch, 2018). The major axis of the **G**-matrix, given by its leading eigenvector, thus identifies the direction in multivariate trait space with the greatest additive genetic variance and hence the greatest potential response to selection (Schluter, 1996). Altogether, the **G**-matrix describes both the capacity of populations to respond to selection and the distribution of this response across traits in the short-term, with broad implications for adaptation, medicine, conservation and evolutionary rescue.

Over longer timescales, the **G**-matrix can itself evolve (Lande, 1984; Steppan et al., 2002; Doroszuk et al., 2008; Arnold et al., 2008; Roff and Fairbairn, 2012; Dugand et al., 2021). Mutation introduces new genetic variance and covariance, as summarised by the **M**-matrix, and can thereby shape the **G**-matrix (Box 1). Recombination reshuffles allelic combinations and erodes covariance generated by linkage disequilibrium (Lande, 1984). Genetic drift causes stochastic changes in allele frequencies and, in expectation, reduces segregating variation (Bürger and Lande, 1994). Selection can also change the **G**-matrix by favouring alleles, or combinations of alleles, that contribute to variation along some trait combinations rather than others (Roff, 2000). Models have examined this process under stabilising selection around a multidimensional optimum (Jones et al., 2003; Milocco and Salazar-Ciudad, 2022; Petit et al., 2023), directional selection towards a moving optimum (Jones et al., 2004), fluctuating selection (Revell, 2007; do O and Whitlock, 2023), and local adaptation with dispersal (Guillaume and Whitlock, 2007). These models show that selection can favour genetic covariance that aligns with the fitness landscape. For example, correlational selection – selection favouring particular combinations of trait values (Lande and Arnold, 1983; Phillips and Arnold, 1989) – can orient the **G**-matrix towards the axis favoured by selection (Jones et al., 2003). The persistence of this genetic covariance however depends on its genetic basis. Covariance caused by pleiotropic alleles is transmitted with those alleles, whereas covariance caused by linkage disequilibrium among recombining loci is eroded by segregation and recombination, although physical linkage can slow this erosion (Lande, 1984; Chebib and Guillaume, 2021).

Most models of **G**-matrix evolution assume many freely recombining additive loci, often with pleiotropic effects. They therefore say little about how the evolution of recombination suppression affects the **G**-matrix. This leaves the effect of supergenes on **G**-matrix evolution understudied. Supergenes are genomic regions in which multiple functional genetics elements are inherited together and jointly control alternative complex phenotypes, such as mimetic wing patterns in butterflies, social organisation in ants, and correlated plumage and behavioural morphs in white-throated sparrows (Schwander et al., 2014; Villoutreix et al., 2021; Saltz et al., 2017; Gutiérrez-Valencia et al., 2021; Berdan et al., 2023). They often involve reduced recombination, including recombination suppression caused by chromosomal inversions, which preserves associations among alleles. Theory has examined the conditions under which inversions evolve after capturing pre-existing alleles with prescribed effects on fitness, often at two or a few diallelic loci (e.g. Kirkpatrick and Barton, 2006; Feder and Nosil, 2009; Charlesworth and Barton, 2018; Mackintosh et al., 2024). These models leave the connection to traits, and therefore to genetic variance in traits, implicit. Far fewer models make this connection explicit by allowing inversions to evolve alongside polygenic traits. In a two-patch model of local adaptation with gene flow, Schaal et al. (2022) showed that inversions can capture standing genetic variation and accumulate small-effect alleles, thereby concentrating a disproportionate share of the additive genetic variance underlying divergence between locally adapted populations within inverted regions. Whether genetic variance similarly concentrates within inversions when disruptive selection acts on multiple traits and favours alternative multi-trait phenotypes remains unclear. Nor is it known how such concentration affects the structure and stability of the **G**-matrix.

Supergenes may also affect genetic (co)variances through dominance. Supergenes are frequently associated with dominant or recessive effects on phenotype (Llaurens et al., 2017). By contrast, most models of **G**-matrix evolution assume purely additive gene action (directly on traits, e.g. Jones et al., 2003, 2004; Guillaume and Whitlock, 2007; Revell, 2007; do O and Whitlock, 2023; or on developmental parameters leading to traits, e.g. Milocco and Salazar-Ciudad, 2022; Petit et al., 2023). Yet dominance can evolve when selection favours discrete phenotypic types but sexual reproduction continually produces intermediate heterozygotes (Otto and Bourguet, 1999). Previous models have examined the evolution of dominance at maintained polymorphisms (e.g. Van Dooren, 1999; Otto and Bourguet, 1999; Doorn and Dieckmann, 2006; Peischl and Schneider, 2010; Spencer and Priest, 2016; Lesaffre et al., 2024, 2025; Flintham, 2025), but not within recombination-suppressed supergenes in relation to additive genetic covariances among multiple traits (though see Lesaffre et al., 2025). Because supergenes are often maintained as balanced polymorphisms that generate recurrent heterozygotes, they may provide particularly favourable conditions for the evolution of dominance. This could alter how much genetic variation remains additive, and therefore how much of the total genetic variance–covariance structure is captured by the **G**-matrix.

Here, we investigate how supergene evolution affects the structure and stability of the **G**-matrix. We consider the joint evolution of two traits that mediate competition for resources and are encoded by multiple pleiotropic loci. By varying the distribution of resources, we generate stabilising selection, disruptive selection that favours diversification only among correlated combinations of trait values, or disruptive selection that favours diversification across both dimensions of trait space. We ask when inversions that suppress recombination among trait-affecting alleles evolve and favour the emergence of supergenes. We also allow dominance to evolve at polymorphic sites. We quantify how supergenes affect heritability, additive genetic correlations, and the temporal stability of the **G**-matrix, and how dominance changes the relationship between additive and total genetic vari-ance–covariance matrices (**G** and **G**_T_-matrices in Box 1, respectively).

## 2 Model

### 2.1 Life cycle, traits, and selection

#### 2.1.1 Life-cycle events and competition for resources

We consider a monoecious population of constant size *N*with non-overlapping generations. Individuals go through the following life-cycle events at each generation: (1) adults compete for resources; (2) each adult produces a large number of gametes according to its fecundity, which is determined by the total amount of resources acquired, and then dies; (3) gametes fuse at random to form zygotes; (4) zygotes compete uniformly for the *N* breeding spots of the next generation.

Each individual *i* ∈ {1,…, *N*} expresses two quantitative traits, denoted 1 and 2, with values *z*_1,*i*_ and *z*_2,*i*_ that we collect in a vector **z***_i_ =* (*z*_1,*i*_, *z*_2,*i*_) and that mediate resource competition. Specifically, we assume that these traits determine how individuals extract resources from the environment through a matching mechanism between resource properties and individual traits (Levene, 1953; Slatkin, 1980; Doebeli, 1996; Day, 2000; Schmid et al., 2024, and references therein). For example, in nectar-feeding birds, beak length may determine access to nectar in flowers differing in corolla-tube depth, whereas beak curvature may determine extraction efficiency from flowers differing in corolla-tube curvature.

We assume four discrete resource types distributed in a two-dimensional property space (fig. 1A). For example, the two properties could be corolla-tube depth and curvature, with the four resource types corresponding to deep or shallow flowers with strongly or weakly curved tubes. We vary two key features of the resource distribution. First, *δ*_u_ controls the amount of variation in each resource property. Specifically, each resource property takes one of two values, −*δ*_u_ or *δ*_u_, and therefore has variance 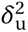. Second, *ρ* changes the relative abundances of the four possible property combinations and thereby controls the correlation between the two resource properties. When *Q =* 0, all four combinations are equally abundant. As *Q* increases, combinations in which the two properties have the same sign become more abundant, e.g. deep flowers with strongly curved tubes and shallow flowers with weakly curved tubes become more common. When *ρ =* 1, only these two positively associated combinations remain. We hold niche breadth fixed (i.e. the rate at which consumption efficiency declines as trait values depart from the corresponding resource properties) and use it to set the scale of trait and resource-property values. Appendix A gives more information regarding resource distribution and competition, and how resource acquisition determines fecundity.

**Figure 1:**
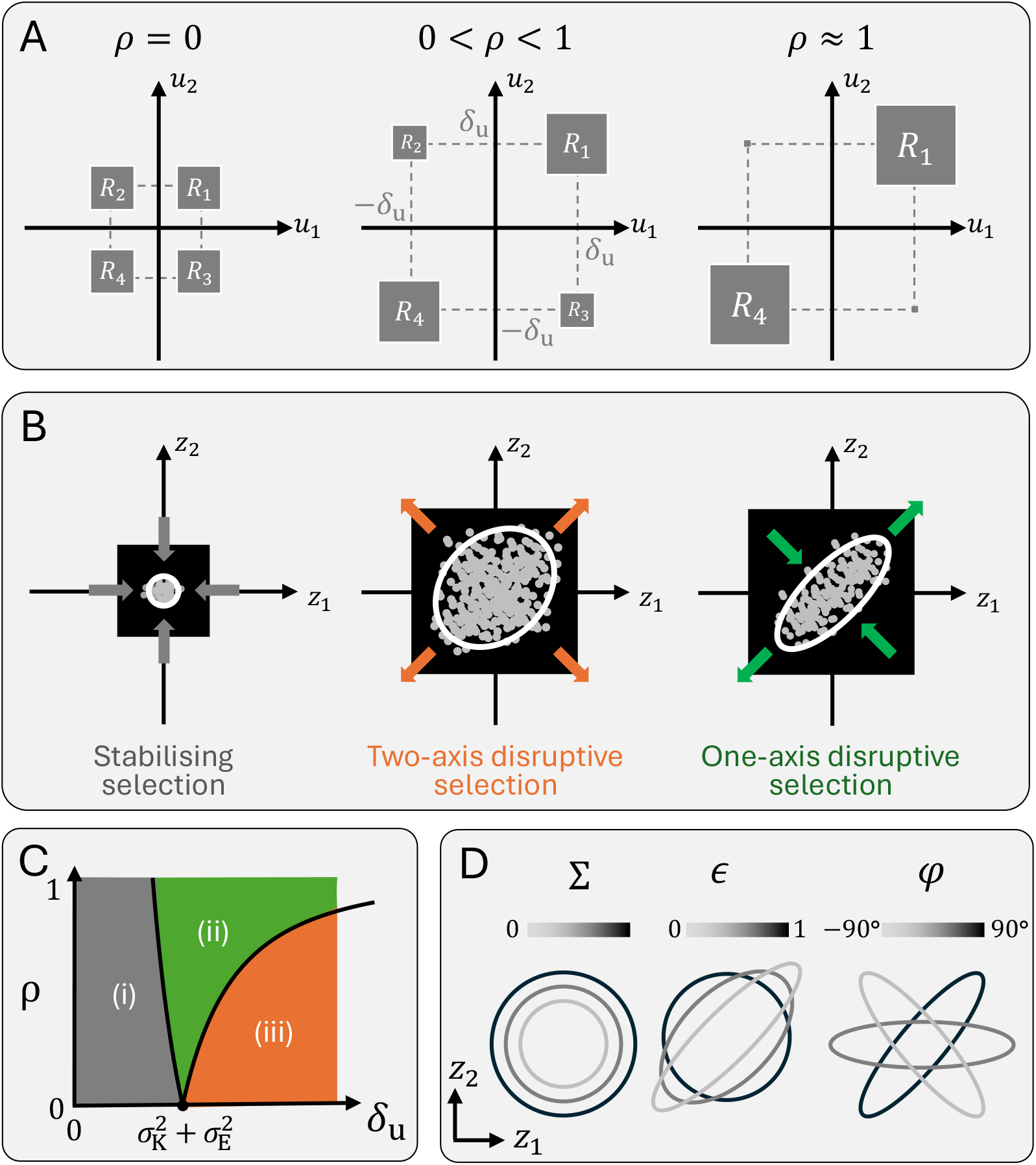
Resource and trait distribution, selection regimes and the elliptical representation of the G-matrix. **A.** Distribution of the four resource types across the two-dimensional property space. The amount of each resource, *R_j_*, determines the correlation *ρ* among the two resource properties in the environment (Appendix A for details). **B.** Examples of phenotype distribution for polygenic ecological traits influencing the consumption of resources in A. Black squares represent the boundaries of resource distribution in A (*±δ*_u_). Arrows indicate the directions in which selection is stabilising (inwards arrows) or disruptive (outward arrows). **C.** Selection regimes according to resource properties’ divergence and correlation (see Appendix B for derivation). **D.** Elliptical representation of a symmetric 2 × 2 matrix (e.g. **P**, **G**_T_, or **G**) based on its eigenstructure described in section 2.3.2: matrix size ( ∑, left), shape (*ε*, middle), and orientation (*φ*, right). Examples illustrate low (light gray), intermediate (gray), and high (black) values of the focal parameter while the remaining parameters are held constant.

#### 2.1.2 Selection on traits: stabilising, disruptive, correlational

Because the two traits determine how efficiently individuals acquire resources, the distribution of resource properties determines selection on these traits (fig. 1B). In particular, *δ*_u_ and *ρ* determine whether selection is stabilising or disruptive and the strength of correlational selection (Lande and Arnold, 1983; Phillips and Arnold, 1989; Débarre et al., 2014). In Appendix B, we characterise selection by assuming individuals reproduce clonally, and their genetic trait values change only through mutations of small effect, as in a continuum-of-alleles model. Each trait also includes an environmental component with variance 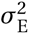, with environmental effects uncorrelated between traits.

The analysis identifies three selection regimes depending on resource-property correlation, *ρ*, resource-property variation, 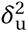, and environmental variance, 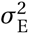 (fig. 1C). We describe selection along two perpendicular axes of trait space: *z*_1_ *= z*_2_, along which the two traits change in the same direction, and *z*_1_ *=* −*z*_2_, along which they change in opposite directions.

**(i) Stabilising selection.** When resource-property variation is sufficiently small relative to niche breadth, such that 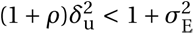, selection is stabilising around the intermediate multivariate optimum (0, 0). In this regime, individuals can efficiently exploit resources with different properties, so intermediate phenotypes are favoured over specialised phenotypes. Correlational selection is nevertheless present when *ρ >* 0: selection penalises phenotypes in which the two traits deviate in opposite directions (*z*_1_ *=* −*z*_2_) more strongly than phenotypes in which they deviate in the same direction (*z*_1_ *= z*_2_).
**(ii) One-axis disruptive selection.** When resource-property variation is intermediate or resource properties are strongly correlated, such that 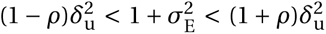, selection is disruptive along *z*_1_ *= z*_2_ but remains stabilising along the perpendicular axis, *z*_1_ *=* −*z*_2_. We refer to this regime as one-axis disruptive selection. Individuals specialising on positively associated combinations of resource properties are favoured, whereas individuals specialising on negatively associated combinations are not. Diversification is therefore favoured only when the two traits vary genetically in the same direction.
**(iii) Two-axis disruptive selection.** When resource-property variation is sufficiently large, such that 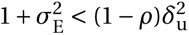, selection is disruptive along both *z*_1_ *= z*_2_ and *z*_1_ *=* −*z*_2_. We refer to this regime as two-axis disruptive selection. Competition and trade-offs in resource use are then sufficiently strong to favour diversification whether the two traits vary in the same or in opposite directions.

These analytical conditions are represented in fig. 1D for different values of resource-property variation, δ_u_, and resource-property correlation, *ρ*; see fig. S1 for confirmation from individual-based simulations.

#### 2.1.3 A polygenic basis to the traits

We are interested in how traits evolve under these different selection regimes when they are encoded by many loci, and how selection in turn reshapes their genetic architecture, in particular through the evolution of supergenes and their effects on the **G**-matrix. We specify here the genetic basis of the two traits.

Each diploid individual carries two homologous chromosomes with *L* pleiotropic loci contributing to the traits. The genotype–phenotype map is additive across loci, but allows dominance within loci. At each locus, an allele is characterised by two quantities: its allelic effects *a* on the two traits, defined as its contribution to trait value when homozygous, and an “affinity” *b >* 0 which determines its relative contribution to each trait when the individual is heterozygous in allelic effects at that locus. This affinity provides a model of phenotypic dominance (Van Dooren, 1999; for applications of this model to polygenic settings: Doorn and Dieckmann, 2006; Lesaffre et al., 2024; Flintham, 2025). It can be interpreted as an allele-specific regulatory effect, for example the affinity of a cis-regulatory sequence for local transcriptional machinery, so that homologous alleles compete for expression at the same locus (Van Dooren, 1999). For simplicity, we assume the same affinity modulates dominance for both traits.

Let *a*_1,*i kℓ*_ denote the effect of the allele carried by individual *i* at locus *k* (*k*∈ {1,…, *L*}) on homolog *l* ∈ {1, 2} on trait 1, *a*_2,*i kℓ*_ for its effect on trait 2, and *b_i kℓ_ >* 0 for its affinity, then the trait values of individual *i* are given by

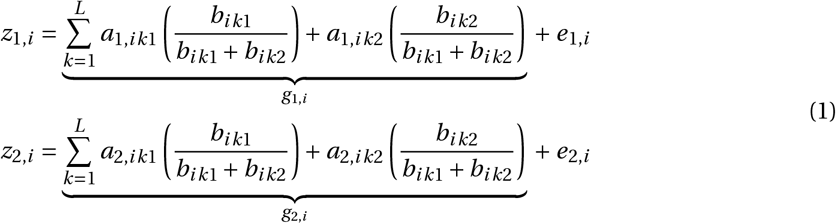

Here, *g*_1,*i*_ and *g*_2,*i*_ are the genetic contributions to traits 1 and 2, respectively. The environmental effects *e*_1,*i*_ and *e*_2,*i*_ are sampled independently with mean zero and variance 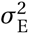. Thus, the **E**-matrix defined in Box 1 is 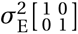. As shown in eq. (1), when the two homologous alleles at a locus have equal affinity (*b_ik_*_1_ *= b_ik_*_2_), they contribute equally to the phenotype. When one allele has higher affinity, its trait effects contribute more strongly, making it dominant relative to the lower-affinity allele.

### 2.2 Evolution of the genetic architecture

We use forward-time, individual-based simulations in SLiM v.5 (Haller et al., 2025) to investigate the evolution of the genetic architecture of the two traits. Each simulation follows the life cycle described above, with selection arising from resource acquisition, inheritance determined by the multilocus genotype–phenotype map, and mutations affecting allelic effects, dominance affinities and structural variants, as described below. Simulation code can be accessed at https://github.com/vsudbrack/SupergenesGmatrix and details regarding the simulation can be found in Appendix C.

We initialise the population without genetic variation in either trait. Wild-type alleles have no effect on the traits, such that *a*_1,*ikℓ*_ *= a*_2,*ikℓ*_ *=* 0, and all alleles have affinity *b_ikℓ_ =* 1. Alleles are therefore initially codominant at all loci. We also assume an initially uniform recombination map, with adjacent loci recombining at rate *r*. See Table 1 for a list of parameters and default values.

**Table 1:** Parameters and default values in the simulations.

| Symbol | Description | Default value |
| --- | --- | --- |
| **Population, traits and resources** |  |  |
| $N$ | Population size | 1'000 |
| $N_R$ | Number of resources available for the population | 4 |
| $\sigma_E^2$ | Variance in environmental effects for each trait | 0.01 |
| $\sigma_K^2$ | Niche breadth | 1* |
| **Genetic architecture** |  |  |
| $L$ | Number of loci | 50 |
| $\mu$ | Allelic mutation rate, per locus per gamete | 0.0002 |
| $\sigma_M^2$ | Variance of mutational increments in allelic effects $a$ | 0.005 |
| $\sigma_B^2$ | Variance of mutational increments in affinity modifier $b$ | 0.05 |
| $r$ | Recombination rate between adjacent loci in homokaryotypes for structural variants | 0.04 |
| **Chromosomal inversions** |  |  |
| $\mu_I$ | Structural mutation rate generating chromosomal inversions, per gamete | $10^{-5}$ |
| $\bar{L}_I$ | Mean inverted stratum length, in number of loci | 5 |

\* In this scaling, we measure the variance between resource properties, $\delta_u^2$ , in units of niche breadth.

#### 2.2.1 Evolution of allelic effects and dominance

Alleles mutate at rate *µ* per locus per gamete. A mutation can alter the effects of an allele on both traits (*a*) and its affinity (*b*).

##### Allelic effects

Allelic effects evolve according to the continuum-of-alleles model used in previous models of **G**-matrix evolution (Jones et al., 2003, 2004; Guillaume and Whitlock, 2007; Chebib and Guillaume, 2021). At each mutation, changes in *a*_1_ and *a*_2_ are drawn from a bivariate normal distribution with mean zero and variance–covariance matrix 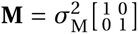, where 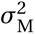 determines the mutation step size. We choose 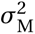 such that the expected genetic variance introduced by mutation each generation satisfies 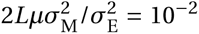, within the range of empirical estimates of mutational heritability (Lynch, 1988; Jones et al., 2003).

We bound allelic effects between −*δ*_u_ /10 and *δ*_u_ /10. Because each resource property takes the value *δ*_u_ or *δ*_u_ (Table 2), reaching a perfect match to a resource-property value requires ten allelic contributions of maximum magnitude in the same direction. For example, ten contributions of δ_u_/10 are required to produce a trait value of *δ*_u_. This constraint prevents a single large-effect locus from generating alternative phenotypic morphs.

##### Dominance through affinity effects

The same mutation can also alter the affinity of an allele. The change in *b* is drawn from a normal distribution with mean zero and variance 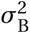. Affinities are constrained to remain positive, ensuring that the relative contributions of homologous alleles in eq. (1) are always defined. Because the same affinity weights both pleiotropic effects of an allele, dominance evolves jointly for the two traits.

#### 2.2.2 Evolution of recombination via inversions and supergenes

We allow the recombination map to evolve through chromosomal inversions. An inversion reverses the orientation of a chromosome segment relative to the ancestral arrangement. Recombination within the inverted segment occurs between chromosomes carrying the same arrangement but is fully suppressed in inversion heterokaryotypes, following Schaal et al. (2022). We thus ignore double crossovers and gene conversion within inversion heterokaryotypes.

Inversions arise at rate *µ*_I_ per gamete. Their length, measured by the number of loci spanned, is drawn from a Poisson distribution with mean *L̅*_I_ (ensuring a minimum length of 2 loci per stratum, sec. C.5), and their position is drawn randomly along the chromosome. When a new inversion overlaps an existing inversion, the overlapping inversions form a single expanded structural variant that recombines only in homokaryotypes.

We say an inversion is segregating when its population frequency lies between 10% and 90%. For each segregating inversion *j*, we estimate its effects on the two traits by regressing individual phenotypes on inversion dosage and heterozygosity: **z***_i_ ∽* ***µ****_j_ +****α****_j_ x_ij_ +****β****_j_ δ_ij_*, where *x_ij_ =* 0, 1 or 2 is the number of copies of inversion *j* carried by individual *i*, and *δ_ij_ =* 0, 1 or 0 when the individual is a standard homozygote, heterokaryotype or inversion homozygote, respectively. The vector ***µ****_j_ =* (*µ*_1,*j*_, *µ*_2,*j*_) gives the average trait values of standard (non-inverted) homozygotes, ***α****_j_ =* (*α*_1,*j*_, *α*_2,*j*_) gives the average change in the two traits per copy of the inversion, and ***β****_j_ =* (*β*_1,*j*_, *β*_2,*j*_) gives the deviation of the heterokaryotype from this additive expectation.

We define a supergene as a segregating inversion with an estimated effect on each trait at least as large as the maximum effect of a single allele: *|α*_1,*j*_ | ≥ *δ*_u_/10 and *|α*_2,*j*_ | ≥ *δ*_u_/10. This definition identifies inversions containing linked genetic variation that jointly controls both traits, following Thompson and Jiggins (2014).

### 2.3 Estimating and representing the G-matrix

#### 2.3.1 A common-garden experiment to obtain offspring–midparent regressions

Every 100 generations, we estimate the additive genetic variance–covariance **G**-matrix using a simulated common-garden experiment. We generate 10*N* offspring from parents sampled randomly from the focal population and mated without selection. This larger sample improves the precision of the offspring–midparent regressions. Once the **G**-matrix has been estimated, the common-garden population is discarded and the focal simulation continues.

Each common-garden offspring *i* has trait values **z***_i_*, given by eq. (1), and corresponding midparent trait values 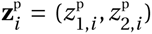, calculated as the mean trait values of its two parents. We estimate the narrow-sense heritabilities of traits 1 and 2 from the slopes of the corresponding offspring–midparent regressions:

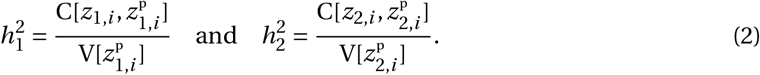

Here and below, variances and covariances are calculated across common-garden offspring *i* and their corresponding midparent values (p. 146 and eqs. 9.4 and 10.1 in Falconer and Mackay, 1997).

Each cross-trait offspring–midparent covariance estimates half the additive genetic covariance between the two traits. We therefore estimate their additive genetic correlation as

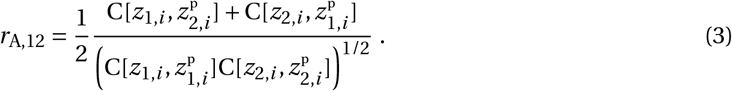

This estimates the correlation between breeding values for the two traits (eq. 19.3 of Falconer and Mackay, 1997).

We then estimate the elements of the **G**-matrix defined in eq. (1.4) of Box 1. The additive genetic variances are

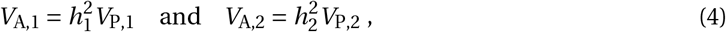

where *V*_P,1_ and *V*_P,2_ are the phenotypic variances of traits 1 and 2 in the population. The additive genetic covariance is

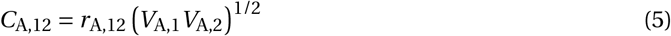

(p. 314 of Falconer and Mackay, 1997).

#### 2.3.2 Elliptical representation of the G-matrix

For two traits, the structure of the **G**-matrix can be represented as an ellipse (fig. 1D; Jones et al., 2003). The eigenvalues of **G** are

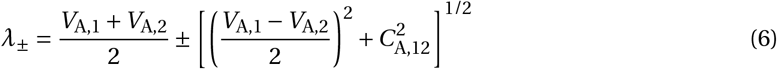

where λ*_+_* ≥ λ_−_. Following Jones et al. (2003), we summarise the **G**-matrix with three quantities. **(i) Its size**, ∑ *=* λ*_+_* +λ_−_, which equals the total additive genetic variance across the two traits (fig. 1D, left). **(ii) Its shape**, *ε =*λ_ /λ*_+_*, which measures how evenly additive genetic variance is distributed across trait combinations. Values of *ε* close to 1 indicate that additive genetic variance is distributed similarly in all directions of trait space, whereas values close to 0 indicate that it is concentrated along one principal axis (fig. 1D, centre). and **(iii) Its orientation**, *φ =* arctan *C*_A,12_ (λ*_+_* - *V*_A,2_), which gives the angle of the major axis relative to the trait-1 axis (−90° *< φ* ≤ 90°). For example, *φ =* 45° indicates that this variance is concentrated along combinations in which both traits change in the same direction, whereas *φ =* 45° indicates that it is concentrated along combinations in which the traits change in opposite directions (fig. 1D, right). When λ*_+_ =* λ−, the **G**-matrix is circular and its orientation is undefined.

This parameterisation facilitates comparisons among matrices without requiring separate comparisons of their individual elements (e.g. fig. 4 in Arnold et al., 2008). The same representation can be applied to other variance–covariance matrices, including the **P**- and **G**_T_-matrices.

## 3 Results

### 3.1 The selection regime determines the structure of the G-matrix

We first examine the structure of the **G**-matrix before allowing inversions or genetic dominance to evolve. During this baseline phase, inversion mutations are absent and allelic effects are additive (*µ*_I_ *=* 0 and 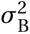 *=* 0; 0 ≤ *t* ≤ 100*’* 000 phase A in fig. 2). We quantify the shape (ε), size (∑) and orientation (*φ*) of **G** by averaging these parameters over 80*’*000 ≤ *t* ≤ 100*’*000 across different resource distributions (left column of fig. 3). This provides a baseline against which we can evaluate the subsequent effects of inversion evolution.

**Figure 2:**
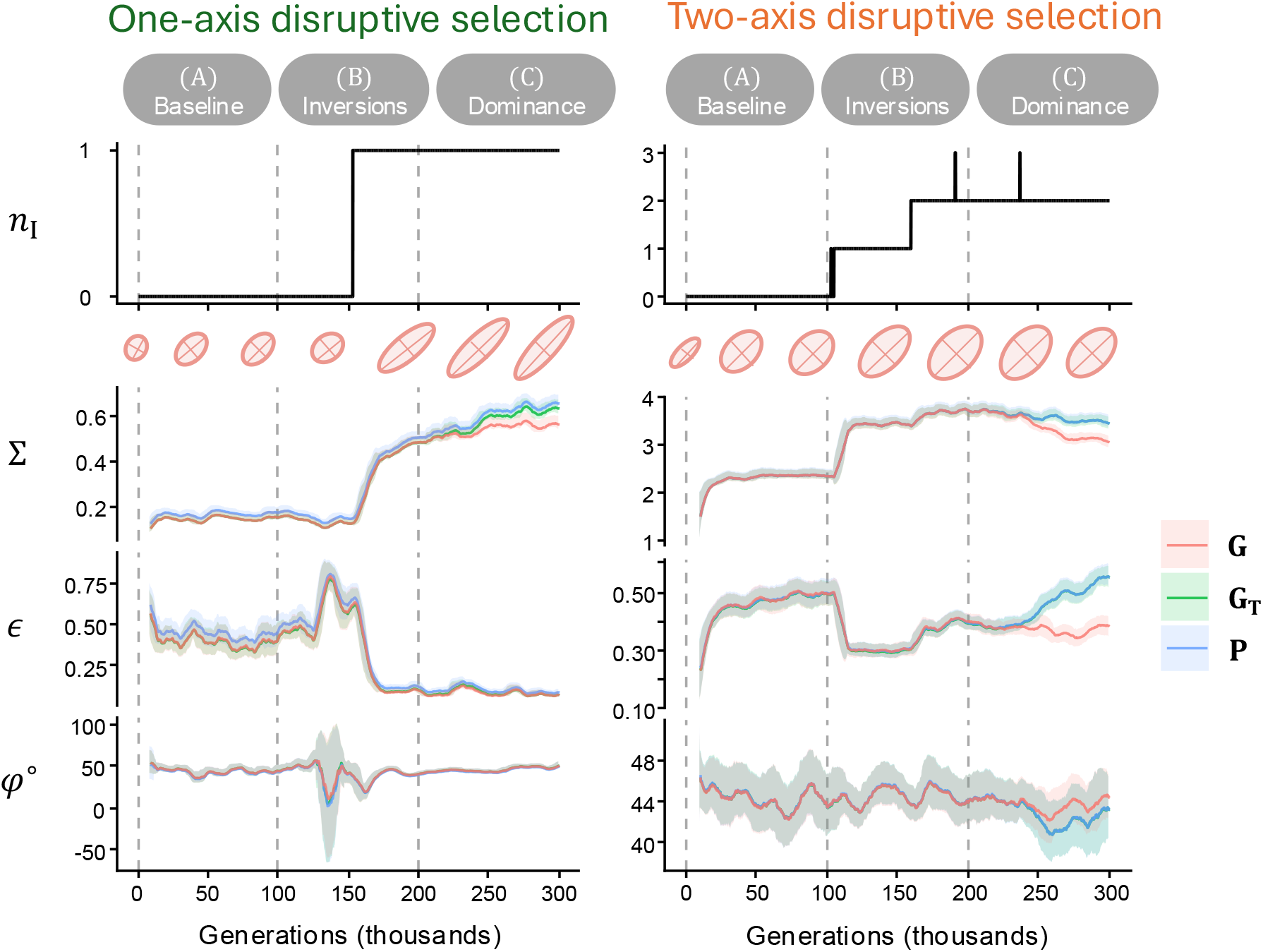
Examples of dynamics of the P-, G_T_- and G-matrices. Evolutionary dynamics under one-axis (left column, *ρ =* 0.1, *δ* _u_ *=* 1.2) and two-axis disruptive selection (right column, *ρ =* 0.2, *δ* _u_ *=* 1.75) in three phases (A: under free recombination and additive effects; B: when inversions can evolve; C: when dominance can evolve). Top : dynamics of the number of chromosomal inversions segregating in the population; here one eventually segregates under one-axis disruptive selection, while two segregate under two-axis disruptive selection. Bottom: dynamics of the shape *ε*, size ∑, and angle *φ* (in degrees) of the phenotypic (**P**; in blue), total genetic (**G**_T_; in green) and additive genetic (**G**; in red) variance-covariance matrices. Solid lines show values averaged over running windows of 10*^’^*000 generations, with shaded area indicating ±1 standard deviation within each window. We also show snapshots of the **G**-matrix as ellipses, with major and minor axes proportional to 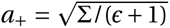and 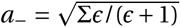 respectively. Fixed parameters: 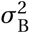 *=* 0, *µ*_I_ *=* 0 during 0 ≤ *t <* 100*^’^*000; *µ*_I_ *=* 10^↑5^, *L̅*_I_ *=* 5 and 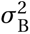 *=* 0 during 100*^’^*000 ≤*t <* 200*^’^*000; and 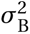 *=* 0.05 and *µ*_I_ *=* 0 during 200*^’^*000 ≤ *t* ≤ 300*^’^*000 (vertical dashed gray lines separate these periods). Other parameters: see table 1.

**Figure 3:**
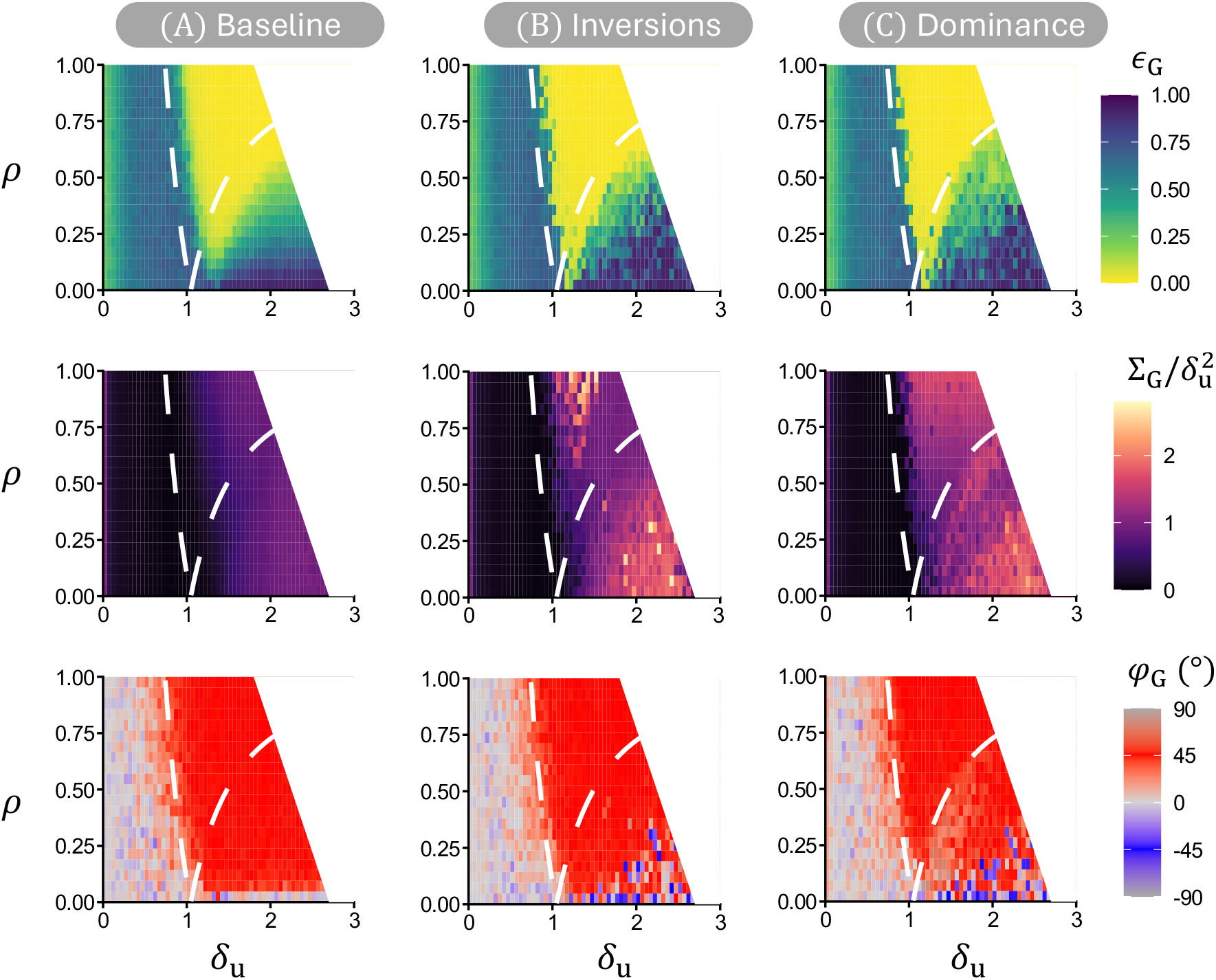
The structure of the G-matrix at the end of each evolutionary phase. The shape, *ε***_G_** (top row), size scaled by the variance of resources, 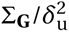 (middle row) and angle, *φ***_G_** (in degrees; bottom row) of the **G**-matrix for various resource distributions in the absence of inversions (80*^’^*000 ≤ *t <* 100*^’^*000; left column), after the evolution of inversions with additive effects (180*^’^*000 ≤ *t <* 200*^’^*000; centre column) and after the evolution of genetic dominance (280*^’^*000 ≤ *t <* 300*^’^*000; right column). White dashed lines delimit the various regimes of selection as in fig. S1 (eq. B8). The white space corresponds to the region of parameter space in which selection is sufficiently strong to generate multimodal phenotypic distributions (see details in fig. S2). Parameters: same as fig. 2.

The structure of **G** is largely determined by the three selection regimes identified in sec. 2.1.2 (dashed lines in fig. 3). We describe these regimes in order of increasing resource divergence, *δ*_u_.

When resource divergence is small, selection is stabilising. In this case, little additive genetic variation is maintained (∑_G_ small; middle-left panel of fig. 3). The **G** matrix remains approximately circular (ε_G_ close to one) and shows no consistent alignment with the direction of resource correlation (variable and weak *φ*_G_), even when correlational selection is present ( *ρ >* 0). Because **G** is small in this regime, its orientation is governed primarily by stochastic fluctuations and new mutations rather than by correlational selection (as observed in Jones et al., 2003).

At intermediate resource divergence, selection becomes disruptive along *z*_1_ *= z*_2_, while remaining stabilising along the perpendicular direction, *z*_1_ *=* −*z*_2_ (i.e. “one-axis disruptive selection” sec. 2.1.2). As a result, the **G** matrix becomes elongated and its major axis aligns with the direction of resource correlation (ε**_G_** close to zero and *φ*_G_≈ 45°; yellow and red regions in the top- and bottom-left panels of fig. 3). This pattern emerges even when the correlation between resource properties *ρ* is weak. By favouring genetic variation along the direction of resource correlation while constraining variation along the perpendicular direction, selection generates strong additive genetic correlations between the two polygenic traits (ε_G_ ≈ 0).

As resource divergence increases further, selection becomes disruptive along both directions of trait space (*z*_1_ *= z*_2_ and *z*_1_ *=* −*z*_2_, i.e. “two-axis disruptive selection” sec. 2.1.2)). Genetic variation is therefore maintained along both trait combinations, and the shape of **G** depends more continuously on the strength of correlational selection (*c*_G_ changes smoothly with *Q*, top left panel of fig. 3). When *ρ >* 0, additive genetic variation remains greater along the direction of resource correlation than along the perpendicular direction (bottom left panel of fig. 3). Correlational selection therefore produces a more elongated **G**-matrix. Only when correlational selection is absent or extremely weak do we see a near-circular **G**-matrix.

At still larger values of *δ*_u_, disruptive selection is sufficiently strong to generate linkage disequilibrium among alleles with similar effects and thereby produce discrete ecological morphs even without inversions. Because inversions are unnecessary for phenotypic morphs to emerge under these conditions, we excluded these simulations from subsequent analyses. Specifically, we excluded cases in which Hartigan’s dip test rejects unimodality at the 1% level at the end of the baseline phase (*t =* 100*’*000). These cases form the white region on the right-hand side of fig. 3 and subsequent figures (see fig. S2).

### 3.2 The form of disruptive selection determines supergene architecture

Following the baseline phase, we allow inversions to arise and spread while allelic effects remain strictly additive (*µ*_I_ *>* 0 and 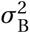 *=* 0 during the B “Inversions” phase, 100*^’^*000 *< t* ≤ 200*^’^*000 in fig. 2). Only under disruptive selection do inversions establish, in which case they persist at intermediate frequencies (fig. 4B). Segregating inversions then often expand through the accumulation of overlapping strata, eventually showing opposite average allelic effects compared to the homologous ancestral segment (e.g. fig. 4A). Most strata capture variants affecting both traits and therefore constitute supergenes according to our operational definition, following Thompson and Jiggins (2014) (yellow region in fig. 4C).

**Figure 4:**
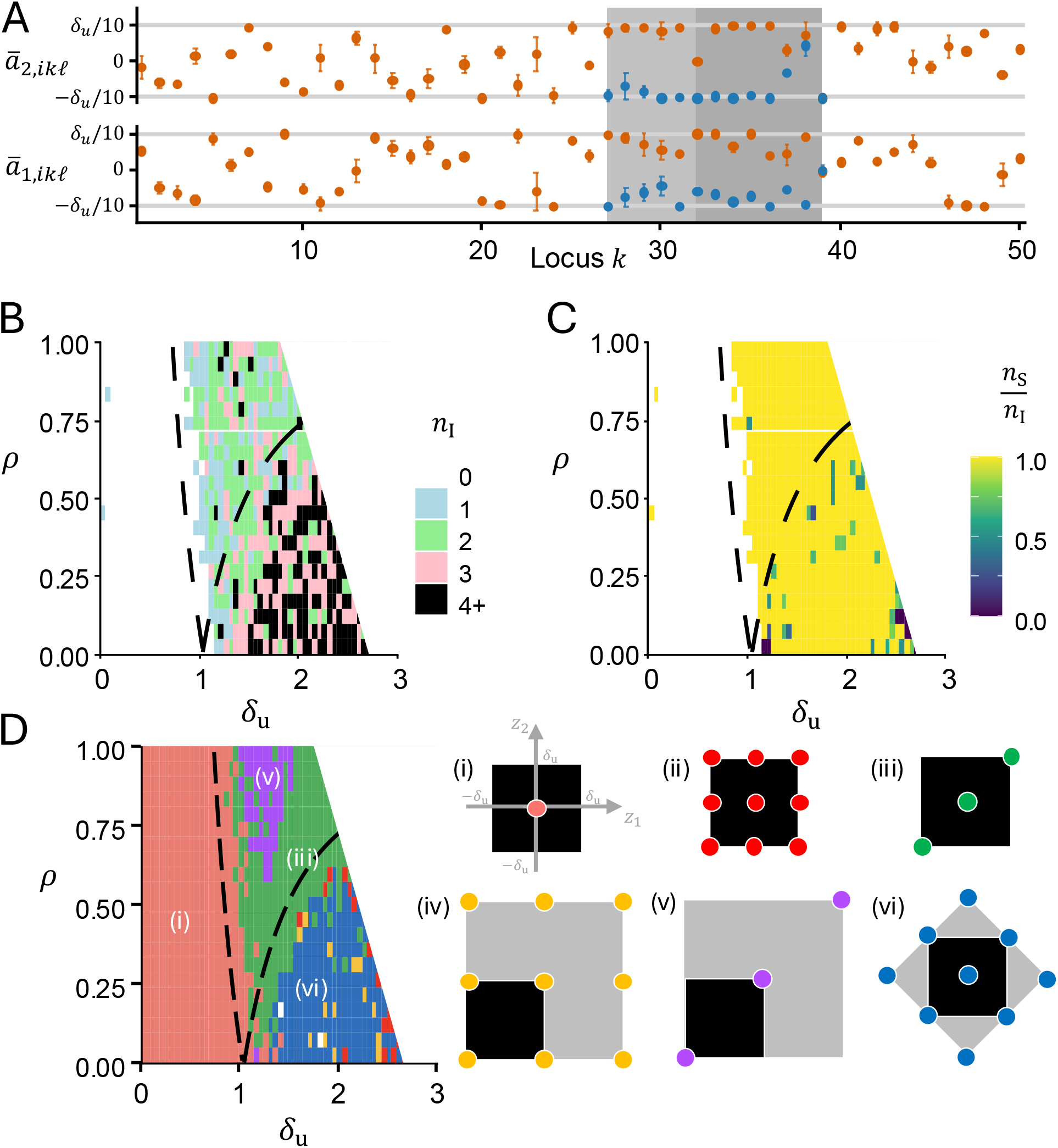
The genetic architecture sustaining niche diversification. **A.** Example of an evolved chromosome. Allele effect on traits *z*_1_ and *z*_2_ at each locus (*k*), averaged across all collinear (red) and inverted (blue) loci (with respect to ancestral order) at *t =* 200*^’^*000. In this simulation, a supergene emerged between loci 26 and 38, composed of two strata (the first between loci 31 and 38, and a second between 26 and 31). Parameters: *ρ =* 0.5, *δ*_u_ *=* 1.5, 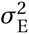 *=* 0.01. **B.** Number *n*_I_ of segregating strata according to selection regime. **C.** Number of supergenes relative to number of inversions, *n*_S_/*n*_I_. Each segregating inverted stratum was classified according to whether it affected one or both traits using phenotype–genotype regressions (section 2.2.2). This shows that most inversions consisted of supergenes by the end of phase B. **D.** Left: The six most common types of polymorphisms that evolve according to selection regime. Right: Diagram of the phenotypic distributions under these six polymorphism type. The black squares represent the four resource property values (*±δ*_u_, *±δ*_u_), and each coloured point represents the mean trait value of a morph. Morphs in the gray regions are “overshooting” the range of resource properties (Van Dooren, 1999). We describe these genetic architectures and derive the approximate equilibrium frequencies of the structural variants underlying these polymorphisms in Appendix D. Examples of phenotypic distribution from the simulations are given in the fig. S4. Fixed parameters: same as fig. 2, observed at generation *t =* 200*^’^*000 (i.e. after 100*^’^*000 generations of inversion evolution, and additive effects).

The resulting suppression of recombination maintains associations among alleles at multiple loci, generating discrete phenotypic morphs specialised on different resources. By the end of the inversion phase, populations converged to six recurrent phenotypic distributions, some involving phenotypic overshooting (when homozygous morphs express trait values beyond the range of resource properties; fig. 4D): (i) monomorphic populations (pink), (ii) two inversions without overshooting (red), (iii) a single supergene without overshooting (green), (iv) two inversions with overshooting (yellow), (v) a single supergene with overshooting (purple), and (vi) two supergenes with overshooting (blue). These configurations differ in the number of independently segregating regions of suppressed recombination and in whether adapted morphs are homozygous or heterozygous for the structural variants. The form of disruptive selection strongly determines which genetic architecture evolves.

Under one-axis disruptive selection, populations almost always evolve a single supergene in one of two configurations: without overshooting (configuration iii, green) or with overshooting (configuration v, purple; fig. 4D; left column of fig. 2). The overshooting configuration occurs mainly when *δ*_u_ is small and *ρ* is large. In this configuration, heterozygotes express one of the adapted phenotypes, whereas homozygotes carrying two copies of the inversion overshoot the range of resource properties (overshooting homozygote occurred at a frequency of only 1/9, limiting its contribution to segregation load, see Appendix D for calculations). The non-overshooting configuration becomes more common as *δ*_u_ increases or *Q* decreases. When resource correlations are weak (*ρ* ≤ ∽ 0.5), heterozygotes can exploit the off-diagonal resources (resources 2 and 3 in fig. 1A), favouring the non-overshooting configuration.

Under two-axis disruptive selection, populations instead typically evolve two independently segregating supergenes or, less commonly, two trait-specific inversions (blue, red and yellow regions in fig. 4D; top right of fig. 2). These architectures generate nine phenotypic morphs and maintain more segregating strata on average (black tiles in fig. 4B). Because only four resource types are available, some morphs are necessarily maladapted. Nevertheless, the two-supergene architecture reduces segregation load by adjusting allele frequencies to limit the production of maladapted homozygotes under random mating (Appendix D for details).

The emergence of inversions also changes the **G**-matrix (fig. 2). One main effect is to increase the size of **G**, ∑**_G_** (compare the left and centre columns of fig. 3). By suppressing recombination, inversions maintain positive linkage disequilibrium among alleles with similar effects, thereby increasing additive genetic variance. As environmental variance remains unchanged, this brings the **G**-matrix closer to the **P**-matrix, thus increasing the heritabilities of both traits (fig. S3). Similar effects have been observed in simulations of inversions controlling a single trait under local adaptation (Schaal et al., 2022). The increase in ∑**_G_** is greatest under phenotypic overshooting because this configuration produces the largest differences in trait values among morphs (compare fig. 4D with the middle row of fig. 3).

Supergenes also affect the shape and orientation of **G**. A single supergene concentrates additive genetic variation along one axis, making **G** more elongated (fig. 2, left column). Under two-axis disruptive selection, the emergence of a second supergene often restores a more isotropic **G** (fig. 2, right column). When correlational selection is weak, the supergene associated with variation along the perpendicular axis of correlational selection can sometimes contribute more additive genetic variance than the supergene associated with the direction of resource correlation. The major axis of **G** can then be oriented at −45° (blue region in the bottom-centre panel of fig. 3).

### 3.3 Disruptive selection promotes coordinated dominance, but most genetic variance remains additive

Our results so far consider that all alleles have equal affinity, which does not evolve (with 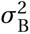 *=* 0 and *b_ikℓ_ =* 1 in eq. 1). Gene action is therefore additive, and the additive and total genetic variance–covariance matrices are identical (**G** *=* **G**_T_ for *t* ≤ 200*^’^*000; fig. 2). We next relax this assumption by allowing affinities to evolve and thus for divergence between **G** and **G**_T_ to occur. Allelic effects also continue to evolve but no new inversions can arise (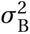 > 0 and *µ*_I_ *=* 0 during the C “Dominance” phase, 200*^’^*000 *< t* ≤300*^’^*000 in fig. 2). We quantify the contribution of dominance to genetic variance as 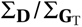, where **D** *=* **G**_T_ −**G** as defined in Box 1 (fig. 5), and regard values of at least 0.1 as substantial (fig. 5).

**Figure 5:**
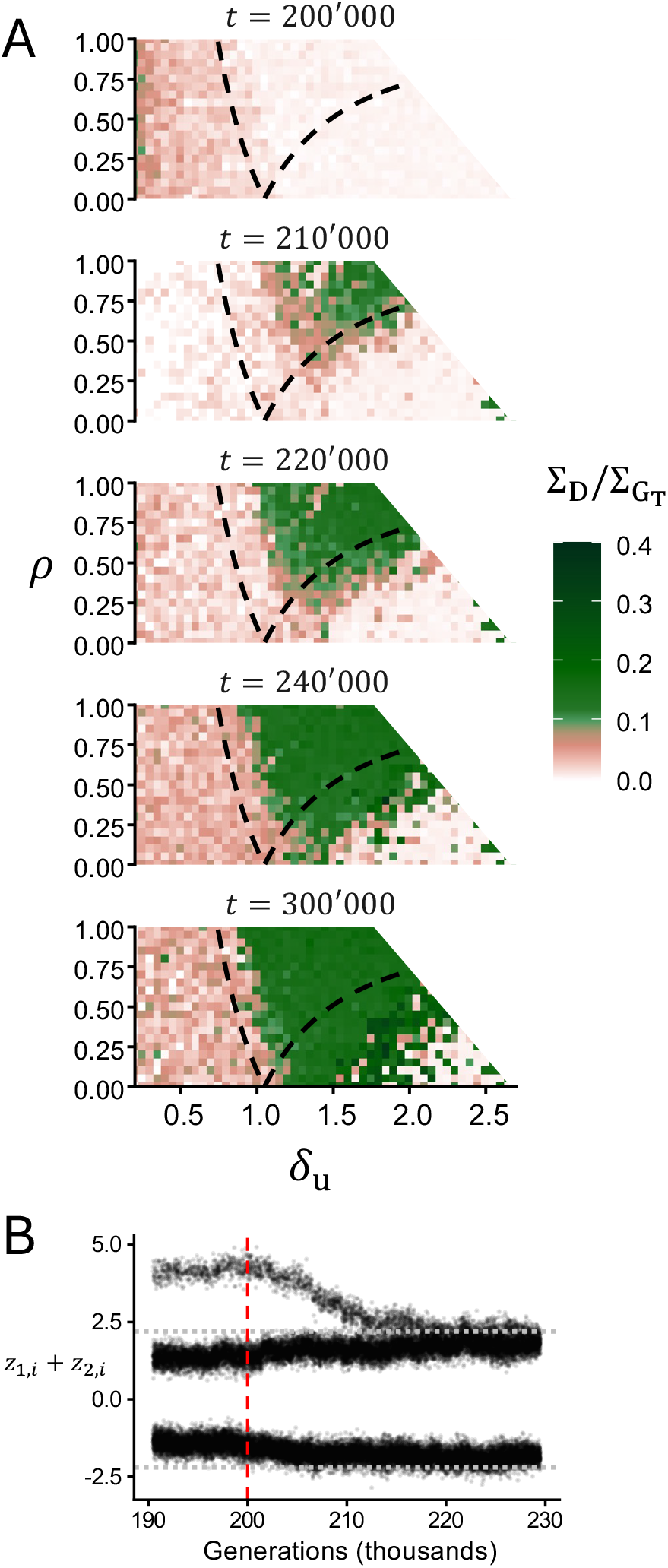
Evolution of non-additive genetic variance when genetic dominance is allowed to evolve. **A.** The relative contribution of dominance to total genetic variance, measured as 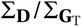, i.e. the ratio between the size of the dominance variance-covariance matrix (**D** *=* **G**_T_ − **G**) and that of the total genetic variance matrix (**G**_T_), at different generations according to selection regime. Values are averaged over 5*^’^*000 generations centred on the time indicated above each panel. Fixed parameters as default in Table 1. **B.** Example of evolution of dominance after the homozygous genotype has overshot under one-axis disruptive selection. As dominance evolves, overshooting collapses so that eventually two phenotypic morphs remain (Supplementary fig. S5 for examples under two-axis disruptive selection). Horizontal dashed lines indicate *±*2*δ*_u_, and vertical red dashed line indicates the onset of dominance evolution. Parameters: *ρ =* 0.8, *δ*_u_ *=* 1.2 and 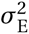 *=* 0.01. Other parameters: see Table 1.

Under stabilising selection, 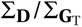 remains low, below 0.1 in most simulations by *t =* 300*^’^*000 (fig. 5). Without mutations on allelic effects, stabilising selection would fix alleles whose homozygous effects produce the optimal phenotype, after which affinity differences would neither be expressed nor selected (Flintham, 2025). Recurrent mutations, however, introduce alleles differing in both their effects and affinities. Mutations that displace the trait from the optimum are selected against less strongly when recessive because their effects are masked in heterozygotes. Recessive mutations therefore segregate for longer, producing a weak apparent increase in dominance variance without generating coordinated dominance across loci.

In contrast, disruptive selection leads to a substantial increase in 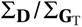 (fig. 5). Because balancing selection maintains inversion polymorphisms under these parameters, heterozygotes are produced recurrently, exposing the phenotypic effects of affinity differences to selection and thereby favouring the evolution of dominance (Otto and Bourguet, 1999). The pace of this increase nevertheless depends on whether selection is one or two-axis disruptive (fig. 5). Under one-axis disruptive selection, populations generally contain a single supergene (configurations iii and v in fig. 4D), and dominance variance accumulates rapidly. Under two-axis disruptive selection, populations generally contain two independently segregating supergenes or inversions (configurations ii, iv and vi), and dominance variance accumulates more slowly. The slower accumulation is presumably because the fitness effect of a change in dominance at one inversion depends on the genotype at the other. The same affinity change can therefore improve resource matching in some genetic backgrounds but not in others, weakening the net selection for coordinated dominance. Interestingly, when *ρ* is small, the two supergenes in configuration (vi) sometimes become trait-specific inversions when dominance evolves (blue region in fig. 4D and fig. S5). In other words, one inversion comes to affect only *z*_1_ and the other only *z*_2_ through joint changes in affinities and allelic effects. Their dominance relationships can then combine independently to generate the four optimal trait combinations matching the four resources (fig. S6). The inversions therefore persist, but they are no longer supergenes under our definition.

More generally, whenever dominance variance becomes substantial, dominance is coordinated across linked loci, i.e. alleles within the same inversion and contributing to the same phenotypic morph tend to acquire higher affinities than their alternatives across multiple loci. Their effects are therefore expressed together in heterozygotes, moving heterozygote phenotypes towards one of the homozygous morphs. This coordination generates a consistent dominant–recessive relationship between chromosomal arrangements across loci.

The effects of dominance evolution on the **G**-matrix are limited, with most genetic variance remaining additive at the population level. Even where coordinated dominance becomes strong, dominance variance commonly accounts for only approximately one-quarter of the total genetic variance summed across the two traits (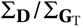 ≈0.25; fig. 5), consistent with single-locus theory showing that strong dominance can coexist with predominantly additive genetic variance (e.g. Hill et al., 2008). The size, shape and orientation of **G** generally change little after the evolution of dominance (centre and right columns of fig. 3). The main exception is a decline in the high ∑_G_ associated with overshooting configuration (v) at small *δ*_u_ and large *ρ*. In this configuration, affinity evolution makes the overshooting allele recessive, and consequently, homozygotes carrying two copies of the arrangement no longer express trait values beyond the range of resource properties, while heterozygotes still express a phenotype matching one resource. The smaller phenotypic differences among genotypes reduce ∑**_G_**.

### 3.4 The effects of inversions on the stability of G depend on the form of disruptive selection

Finally, we ask whether segregating inversions are associated with short-term fluctuations around the mean structure of the **G**-matrix (shaded area around the means in fig. 2). Following Jones et al. (2003), we measure the standard deviations of shape (∑**_G_**) and orientation (*φ***_G_**), and the coefficient of variation of size (∑**_G_**), within 1*^’^*000-generation windows containing 10 estimates sampled every 100 generations. We use the coefficient of variation because the scale of ∑**_G_** depends on trait units and its mean differs among simulations (Jones et al., 2003). It therefore measures fluctuations relative to mean size rather than absolute fluctuations. We classify a window as containing inversions when at least one inversion remains between 10% and 90% frequency at all 10 sampled times. Windows without inversions contain no inversion in this frequency range at any sampled time, and windows in which an inversion entered or left this range are excluded from the analysis. We average the fluctuation metrics across qualifying windows within each simulation. We then compare baseline windows from 80*^’^*000 ≤ *t* ≤100*^’^*000, when inversions are absent, with inversion-phase windows from 180*^’^*000 ≤ *t* ≤ 200*^’^*000 that contain inversions (left column of fig. 6; Appendix E). For dominance, we retain simulations in which the mean size of **D** is at least 10% of that of **G**_T_ during 280*^’^*000 ≤ *t* ≤ 300*^’^*000, and compare this interval with the additive phase at 180*^’^*000 ≤ *t* ≤200*^’^*000 (right column of fig. 6). Negative values in fig. 6 thus indicate lower fluctuations in the presence of inversions (left) or after the evolution of dominance (right).

**Figure 6:**
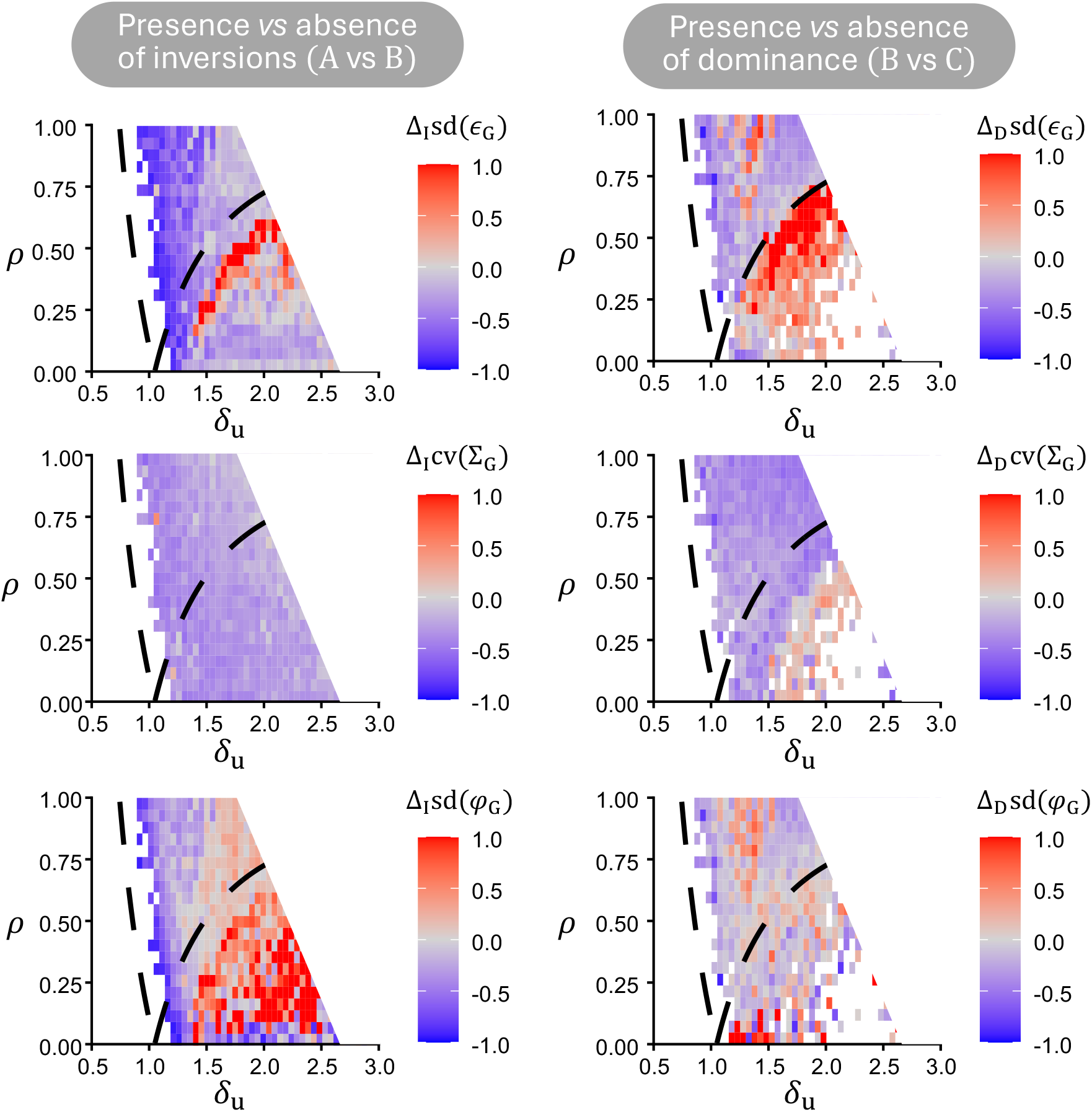
Stability of the G-matrix in the presence versus absence of inversions (left) and of genetic dominance (right). The left column shows the relative change in fluctuations between periods with and without segregating inversions (Δ_I_). The right column shows the corresponding change between phases with evolved dominance and purely additive effects (Δ_D_). Negative values (blue) indicate smaller fluctuations and greater stability; positive values (red) indicate larger fluctuations and lower stability. Rows show fluctuations in shape (*ε***_G_**, top), size (∑**_G_**, middle), and orientation ( *φ***_G_**, bottom). Fluctuations are measured within sliding 1*^’^*000-generation windows using the standard deviation for shape and orientation, and the coefficient of variation for size. The latter measures fluctuations relative to mean size. See Appendix E for calculation and comparison details.

Under one-axis disruptive selection, windows with segregating inversions generally show lower values for all three fluctuation metrics (blue regions in the left column of fig. 6). Established inversion polymorphisms are therefore associated with greater short-term stability of the size, shape and orientation of **G** in this selection regime. This is likely because supergenes reduce the number of genetic components that can fluctuate independently. Without a supergene, drift changes allele frequencies at many loci, while recombination reshuffles their associations, continually changing the distribution of genetic variation across trait space. Within a supergene, these alleles are instead inherited as a single block. Drift and recombination therefore have less scope to alter the size, shape and orientation of **G**.

Under two-axis disruptive selection, segregating inversions remain associated with lower relative fluctuations in ∑**_G_**, but shape and orientation often fluctuate more (red regions in the top- and bottom-left panels of fig. 6). Populations in this regime generally contain two independently segregating regions of recombination suppression, so genetic variance is distributed more evenly between its two principal axes and **G** is more isotropic. As ε**_G_** approaches one, the two axes carry similar amounts of genetic variance, so neither stands out as the main direction of variation. Small changes in this balance may therefore switch the major axis between 45° and −45° (bottom-centre panel of fig. 3). Thus, the greater fluctuations in orientation largely reflect the more isotropic **G** observed when supergenes underlay the traits than when variation is distributed across a polygenic background (sup. fig. S7, which relates the structure of **G** to its temporal fluctuations across selection regimes).

The comparison between the additive and dominance phases shows more heterogeneous and less systematic changes (right column of fig. 6). Relative fluctuations in ∑**_G_** are generally lower after the evolution of dominance, when dominance has an effect. Fluctuations in shape increase across part of the region where selection is two-axis disruptive, whereas orientation shows no clear directional response. Overall, segregating inversions are most consistently associated with lower relative fluctuations in **G** size, whereas their associations with shape and orientation depend on the selection regime. The weaker and less consistent changes after the evolution of dominance agree with our earlier finding that dominance has little effect on the mean structure of **G**.

## 4 Discussion

The **G**-matrix summarises additive genetic variation within a population and can predict the rate and direction of short-term multivariate evolution (Lande, 1979; Lande and Arnold, 1983). It is therefore often treated as a fixed property of a population (Arnold et al., 2008). Over longer timescales, however, mutation, recombination, genetic drift and selection can change its size, shape and orientation (Steppan et al., 2002; Arnold et al., 2008). Much theoretical work on the evolution of the **G**-matrix has considered stabilising selection around a multivariate optimum (Jones et al., 2003; Milocco and Salazar-Ciudad, 2022; Petit et al., 2023). Other models have considered directional selection, fluctuating selection and local adaptation (Jones et al., 2004; Revell, 2007; do O and Whitlock, 2023; Guillaume and Whitlock, 2007), but not how multivariate disruptive selection shapes the **G**-matrix when inversions can evolve to concentrate pleiotropic alleles into supergenes. Our results address this by asking when supergenes evolve under disruptive and correlational selection acting on two coevolving traits, and how their evolution changes the structure and temporal stability of the **G**-matrix.

Our results indicate that the selection regime determines whether supergenes evolve and how genetic variation is organised within the **G**-matrix. Under stabilising selection, little additive genetic variation is maintained, as in previous models of **G**-matrix evolution (Jones et al., 2003; Milocco and Salazar-Ciudad, 2022; Petit et al., 2023), and inversions do not establish. When diversification is favoured along a joint combination of the two traits (i.e. under one-axis disruptive selection), additive genetic variation becomes concentrated along that combination, and populations generally evolve a single supergene controlling both traits. When selection favours diversification in each trait independently (i.e. under two-axis disruptive selection), additive genetic variation is maintained across both traits, and populations may evolve two independently segregating supergenes or, less often, two trait-specific inversions. The form of disruptive selection therefore determines whether genetic variation is captured by a single multi-trait supergene, multiple supergenes or trait-specific inversions, and consequently how additive genetic variation is distributed across trait space (figs. 3 and 4).

The evolution of supergenes in our simulations results from selection against recombination among alleles contributing to alternative phenotypic morphs. Under disruptive selection, recombination breaks down associations among co-adapted alleles and produces intermediate trait combinations that match the environment poorly. Inversions preserve these associations by suppressing recombination in heterokaryotypes and are therefore maintained at intermediate frequencies by balancing selection. The resulting linkage disequilibrium increases additive genetic variance and narrow-sense heritability for both traits (figs. 3 and S3). The concentration of genetic variation within supergenes extends previous models in which variation for a single trait becomes concentrated within fewer loci under local competition (Doorn and Dieckmann, 2006), or within tightly linked genomic regions under local adaptation with gene flow (Yeaman and Whitlock, 2011; Yeaman, 2022; Schaal et al., 2022).

Our results can be related to some of the patterns that have been observed in natural supergenes. First, our finding that supergenes can concentrate genetic variation underlying complex, multi-trait phenotypes is consistent with quantitative-genetic evidence from *Littorina saxatilis*. In this species, linkage groups containing putative inversions contribute disproportionately to additive genetic variation in several shell and foot traits that differ between the Crab and Wave ecotypes (although genetic variation outside these regions also contributes, Koch et al., 2021, 2022). Whether correlational selection favoured the evolution of these inversions in *L. saxatilis* remains unclear, and reconstructing the historical selection pressures is likely to be difficult. Second, our simulations show that a single supergene can generate phenotypic overshooting in inversion homozygotes, whose trait values lie beyond the adaptive range. In the white-throated sparrow, the ZAL2^m^ rearrangement is associated with alternative plumage and behavioural phenotypes (Thomas et al., 2008; Maney et al., 2020). One ZAL2^m^/ZAL2^m^ homozygote that was systematically characterised showed exaggerated plumage, vocalisation and aggression relative to the usual ZAL2/ZAL2^m^ heterokaryotype (Horton et al., 2013). This observation resembles phenotypic overshooting because the homozygote expressed more extreme trait values than the heterokaryotype. Additional ZAL2^m^/ZAL2^m^ homozygotes would be needed to establish whether this genotype consistently produces an exaggerated phenotype, and fitness measurements would be required to determine whether that phenotype lies beyond an adaptive optimum.

Our results further indicate that supergenes maintained under disruptive selection create conditions that favour coordinated dominance, although most genetic variance remains additive. Previous models show that dominance can evolve when balancing selection maintains polymorphism and repeatedly produces heterozygotes, thereby exposing dominance relationships to selection (Van Dooren, 1999; Otto and Bourguet, 1999; Doorn and Dieckmann, 2006). In our simulations, balancing selection similarly maintains supergene polymorphisms and repeatedly produces heterokaryotypes, allowing selection to act on differences in affinity between alternative alleles. Alleles contributing to the same phenotypic morph consequently evolve similar dominance relationships across loci within the same supergene. Nevertheless, dominance variance generally accounts for only approximately one quarter of total genetic variance, even when dominance is strongly coordinated within supergenes (fig. 5). Strong dominance at the level of gene action therefore does not imply that dominance variance predominates at the population level, consistent with single-locus theory (Hill et al., 2008). Accordingly, the size, shape and orientation of the **G**-matrix generally change little after dominance evolves. An exception occurs in populations that evolve phenotypic overshooting while allelic effects are constrained to be additive. When dominance is subsequently allowed to evolve, phenotypic overshooting disappears, reducing phenotypic differences among supergene genotypes and therefore the size of the **G**-matrix (figs. 3 and 5).

By influencing the structure and amount of additive genetic variation, supergenes can facilitate or hamper adaptation, depending on the history of selection and how the environment changes. Our results show that when disruptive selection acts only along the direction of resource correlation, supergenes concentrate additive genetic variation along this direction. Populations should therefore respond more readily if, for example, both resource properties increase together, favouring an increase in both traits. By contrast, if one resource property increases while the other decreases, selection would favour a response along the perpendicular direction, where little additive genetic variation is available (Pavli č ev and Cheverud, 2015). Suppressed recombination may further constrain this response by limiting the formation of new combinations of alleles carried by different chromosomal arrangements (Faria et al., 2019; Roesti et al., 2022). At the population level, responses to selection can involve changes in morph frequencies, changes in trait values within morphs, or both. For example, population mean traits can increase simply because a morph with higher trait values becomes more common, even if the trait values expressed within each morph remain unchanged.

Our estimates of **G** include genetic differences among morphs, so a large population-level matrix does not necessarily imply substantial additive genetic variation within each morph. Given the large trait differences generated by supergenes in our simulations, we expect additive genetic variation within morphs to be small relative to that among morphs. For comparison, most broad-sense genetic variance in fecundity and juvenile size lies among introduced morphs of the predominantly clonal snail *Melanoides tuberculata* (Facon et al., 2008). Whether morphs can evolve independently also depends on genetic covariances between morphs, which are described by cross-morph **B**-matrices. Developed to study genetic covariances between traits expressed in males and females (Lande, 1980), this framework can also be applied to ecological morphs (Roff and Fairbairn, 2011; Wolak et al., 2015). The diagonal elements of **B** give the additive genetic covariance for the same trait expressed in two morphs, whereas its off-diagonal elements give covariances between different traits expressed in the two morphs (Barker et al., 2010). A positive diagonal element would indicate that selection for an increase in a trait in one morph is expected to increase the same trait in the other. A negative diagonal element would instead predict a decrease in the other morph. Together with morph-specific **G**-matrices, these covariances would help establish how independently morphs can respond to selection.

Our results further indicate that segregating inversions can stabilise the **G**-matrix through time. This is especially the case when diversification is favoured along a joint combination of the two traits (one-axis disruptive selection): inversions reduce fluctuations both in the overall amount of additive genetic variation and in its distribution across trait combinations. In this case, a single supergene concentrates much of the genetic variation underlying the alternative morphs within one block. By contrast, when diversification is favoured in each trait independently (two-axis disruptive selection), inversions still stabilise the overall amount of additive genetic variation, but its distribution across trait combinations fluctuates more (fig. 6, left column). This probably occurs because two independently segregating inversions maintain similar amounts of variation along different trait combinations, so small changes in their balance can change which combination contains the most variation.

Of course, our model relies on many simplifications. We highlight two that we think are particularly relevant here. First, we consider only two traits. With more traits, selection could favour several overlapping trait combinations, and genetic variation could become concentrated within one large supergene, distributed among several supergenes or remain polygenic. Second, we assumed that the genotype–phenotype map is additive among loci and excludes epistasis, while dominance evolves only through allele-specific affinity. Epistasis and nonlinear genotype–phenotype maps could generate different patterns of trait covariation and change which supergenes evolve (Milocco and Salazar-Ciudad, 2022; Petit et al., 2023). It also remains to be seen whether coordinated dominance would emerge when dominance arises through other mechanisms, such as thresholds or saturation in metabolic pathways, sensitivity to gene dosage, or interactions in which the protein produced by one allele interferes with that produced by the other (Wilkie, 1994; Bourguet and Raymond, 1998).

## 5 Conclusion

Disruptive selection can favour inversions that preserve associations among alleles contributing to alternative phenotypic morphs, forming supergenes that increase additive genetic variation and heritability and make the **G**-matrix more stable through time. One message from our results is that not all forms of disruptive selection have the same consequences for genetic architecture and heritable variation. When ecology favours diversification along a joint combination of two traits, genetic variation generally concentrates within a single supergene controlling both traits. When ecology instead favours diversification in each trait independently, variation may be spread among several independently segregating inversions. Ecology therefore determines which morphs evolve, as well as whether their genetic basis is packaged within one genomic region or distributed across several, with potentially lasting consequences for the genetic variation available for future evolution.

## Acknowledgements

We thank Aaditya Narasimhan, Ehouarn Le Faou, Ewan Flintham, Isabela do Ó and Mark Kirkpatrick for interesting discussions and feedback on this project. VS thanks the Sachdeva lab for feedback and support during the final stages of this work.

## Conflict of Interest

The authors declare no conflict of interest.

## Box 1

**Multi-trait evolution summarized in variance-covariance matrices**

Quantitative genetics describes the joint variation of continuous traits using variance–covariance matrices (Ch. 8 in Falconer and Mackay, 1997). For two traits, *z*_1_ and *z*_2_, the phenotypic variance–covariance matrix is

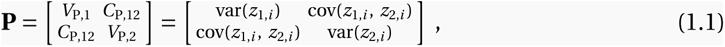

where the (co)variances are computed across individuals (i.e. across *i*). The diagonal elements of the **P**-matrix give the phenotypic variance in each trait, and the off-diagonal elements give the phenotypic covariance between traits.

Phenotypic values can be decomposed into genetic and environmental components, such that *z*_1,*i*_ *= g*_1,*i*_ *+e*_1,*i*_ and *z*_2,*i*_ *= g*_2,*i*_ *+e*_2,*i*_. The total genetic variance–covariance matrix is

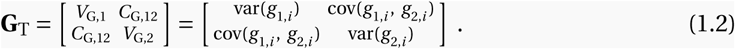

An environmental variance–covariance matrix, **E**, can be defined analogously. If genetic and environmental effects are uncorrelated, these matrices satisfy

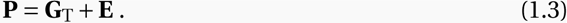

Common-garden experiments control environmental differences among populations or genotypes. When combined with pedigree or genomic information, they can be used to partition phenotypic variation into genetic and environmental components (de Villemereuil et al., 2016).

The genetic component can be further partitioned into additive, dominance and epistatic components: *g*_1,*i*_ *= a*_1,*i*_ *+d*_1,*i*_ *+ε*_1,*i*_ and *g*_2,*i*_ *= a*_2,*i*_ *+d*_2,*i*_ *+ε*_2,*i*_. Here, *a*_1,*i*_ and *a*_2,*i*_ are the breeding values of individual *i*, *d*_1,*i*_ and *d*_2,*i*_ arise from interactions between alleles at the same locus, and *ε*_1,*i*_ and *ε*_2,*i*_ arise from interactions among loci. The additive genetic variance–covariance matrix is

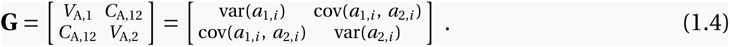

We assume no epistasis, i.e. *ε*_1,*i*_ *= ε*_2,*i*_ *=* 0, so that the difference **D** *=* **G**_T_ - **G** gives the dominance variance–covariance matrix in our model. Estimating the **G**-matrix requires information about relatedness among individuals; here, we use offspring–midparent regressions (section 2.3.1; Wilson et al., 2010).

The structure of the **G**-matrix reflects pleiotropy and linkage disequilibrium among segregating alleles (Lande, 1984). Linkage disequilibrium can arise through physical linkage or selection, including the Bulmer effect (p. 553 in Lynch and Walsh, 1998). The **M**-matrix instead describes mutational input: its elements are the variances and covariances among the trait effects of *de novo* mutations (Lande, 1984; Jones et al., 2007). The off-diagonal elements therefore measure mutational covariance between traits, including covariance generated by pleiotropic mutations. The **M**-matrix can be estimated using mutation-accumulation experiments (Cai et al., 2025).

Together, these matrices connect mutational input, standing phenotypic variation and evolutionary change. The **M**-matrix describes the input of new genetic variance and covariance, the **P**-matrix describes the phenotypic variation exposed to selection, and the **G**-matrix determines the response of trait means to selection (Lande, 1979, 1984; Jones et al., 2007).

## Notes

https://github.com/vsudbrack/SupergenesGmatrix

### Competing Interest Statement

The authors have declared no competing interest.

### Funding

- Swiss National Science Foundation: PCEFP3181243

