## Appendices for "The effects of supergene evolution on the structure and stability of the G-matrix"

for

#### List of appendices

A Resource distribution and competition

B Invasion analysis: directional, quadratic and correlational selection

C Individual based simulations

D Characterization of common trait distributions in the presence of inversions

E Metrics of stability for the **G**-matrix

### A Resource distribution and competition

Here we provide details on our model of resource competition and how that affects fitness.

We consider $N_\mathrm{R} = 4$ resource types available for consumption to all individuals in the population. Resource $j \in \{1,2,3,4\}$ has abundance $R_j$ at the beginning of each generation and two property values, which we collect in the vector $\mathbf{u}_j = (u_{1,j}, u_{2,j})$. Each property takes one of two values, $-\delta_\mathrm{u}$ or $\delta_\mathrm{u}$. The parameter $\delta_\mathrm{u}$ sets the scale of variation in resource property, whereas $\rho \in [0,1]$ determines the association between the two properties through the relative abundances of the four resource types (Table 2).

| $j$ | $u_{1,j}$ | $u_{2,j}$ | $R_j$ |
|---|---|---|---|
| 1 | $\delta_\mathrm{u}$ | $\delta_\mathrm{u}$ | $R(1+\rho)$ |
| 2 | $\delta_\mathrm{u}$ | $-\delta_\mathrm{u}$ | $R(1-\rho)$ |
| 3 | $-\delta_\mathrm{u}$ | $\delta_\mathrm{u}$ | $R(1-\rho)$ |
| 4 | $-\delta_\mathrm{u}$ | $-\delta_\mathrm{u}$ | $R(1+\rho)$ |

**Table 2:** Resource-property values and initial abundances for the four resource types. The parameter $\delta_\mathrm{u}$ sets the scale of resource-property variation. The parameter $\rho$ changes the relative abundances of the four property combinations and, for $\delta_\mathrm{u} > 0$, equals the abundance-weighted correlation between the two properties (fig. 1B).

The total resource abundance is $R_\mathrm{tot} = \sum_{j=1}^{N_\mathrm{R}} R_j = 4R$ for all values of $\rho$. We use the relative abundance $p_j = \frac{R_j}{R_\mathrm{tot}}$ as the weight of resource type $j$. The abundance-weighted means of the two resource properties are

$$\overline{u} = \sum_{j=1}^{N_\mathrm{R}} p_j u_{1,j} = 0 \qquad \text{and} \qquad \overline{v} = \sum_{j=1}^{N_\mathrm{R}} p_j u_{2,j} = 0. \tag{A1}$$

Their variances are

$$\begin{aligned} \sigma^2_{\mathrm{R},1} &= \sum_{j=1}^{N_\mathrm{R}} p_j (u_{1,j} - \overline{u})^2 = \delta_\mathrm{u}^2, \\ \sigma^2_{\mathrm{R},2} &= \sum_{j=1}^{N_\mathrm{R}} p_j (u_{2,j} - \overline{v})^2 = \delta_\mathrm{u}^2. \end{aligned} \tag{A2}$$

Thus, $\sigma^2_{\mathrm{R},1} = \sigma^2_{\mathrm{R},2} = \sigma^2_\mathrm{R} = \delta^2_\mathrm{u}$. Their covariance is

$$\mathrm{cov}_\mathrm{R}(u,v) = \sum_{j=1}^{N_\mathrm{R}} p_j (u_{1,j} - \overline{u})(u_{2,j} - \overline{v}) = \rho\delta_\mathrm{u}^2. \tag{A3}$$

For $\delta_\mathrm{u} > 0$, the abundance-weighted correlation between the resource properties is therefore

$$\mathrm{cor}_\mathrm{R}(u,v) = \frac{\mathrm{cov}_\mathrm{R}(u,v)}{\sqrt{\sigma^2_{\mathrm{R},1}\sigma^2_{\mathrm{R},2}}} = \rho. \tag{A4}$$

When $\rho = 0$, all resource types are equally abundant (Table 2). Increasing $\rho$ increases the abundances of resource types 1 and 4, whose two properties have the same sign, and decreases the abundances of types 2 and 3, whose properties have opposite signs. At $\rho = 1$, only resource types 1 and 4 remain. We restrict the analysis to non-negative values of $\rho$, because a negative association can be obtained by reversing the sign of one resource property.

The efficiency $\kappa_j$ with which individual $i$ consumes resource $j$ is

$$\kappa_j(\mathbf{z}_i) = \exp\left(-\frac{\left(z_{1,i} - u_{1,j}\right)^2}{2\sigma^2_{\mathrm{K},1}} - \frac{\left(z_{2,i} - u_{2,j}\right)^2}{2\sigma^2_{\mathrm{K},2}}\right). \tag{A5}$$

Consumption efficiency therefore declines as either trait departs from the corresponding resource property. The quantities $\sigma^2_{\mathrm{K},1}$ and $\sigma^2_{\mathrm{K},2}$ give the characteristic niche widths along the two ecological dimensions. We assume equal niche widths, such that $\sigma^2_{\mathrm{K},1} = \sigma^2_{\mathrm{K},2} = \sigma^2_{\mathrm{K}}$, and measure trait and resource-property values in units of the common niche width, setting $\sigma^2_{\mathrm{K}} = 1$. Thus, $\delta_{\mathrm{u}}$ is measured relative to $\sigma^2_{\mathrm{K}}$.

Individuals compete for each resource in proportion to their consumption efficiency. The share of resource $j$ acquired by individual $i$ is

$$k_{ij} = \frac{\kappa_j(\mathbf{z}_i)}{\sum_{k=1}^{N} \kappa_j(\mathbf{z}_k)}. \tag{A6}$$

This allocation rule follows from an explicit model of within-generation resource depletion when consumption continues until all resources have been consumed (see Box 1 in Schmid et al., 2024 for a derivation). When all individuals express the same traits, each receives a share $1/N$ of every resource. Otherwise, individuals whose traits better match a resource's properties acquire a larger share of that resource.

Finally, fecundity (i.e. an individual's relative contribution to the population's gamete pool) is proportional to total resource acquisition. Choosing units in which the conversion from acquired resources to offspring is one gives

$$f_i = \sum_{j=1}^{4} R_j k_{ij}. \tag{A7}$$

This formulation assumes that all resource types have the same energetic value per unit and contribute interchangeably to reproduction.

#### B Invasion analysis: directional, quadratic and correlational selection

We investigate the long-term evolution of traits $z_1$ and $z_2$ in a large, effectively infinite population in the absence of genetic constraints. For this analysis, we assume that individuals reproduce clonally. We consider a rare mutant with genetic trait values $\mathbf{g}_\mathrm{m} = (g_{1,\mathrm{m}}, g_{2,\mathrm{m}})$ arising in a resident population that is monomorphic for the genetic trait values $\mathbf{g} = (g_1, g_2)$. Mutational effects are small, such that both $|g_{1,\mathrm{m}} - g_1|$ and $|g_{2,\mathrm{m}} - g_2|$ are small. The realised phenotypes of mutant and resident individuals are $\mathbf{z}_\mathrm{m} = \mathbf{g}_\mathrm{m} + \mathbf{e}$ and $\mathbf{z} = \mathbf{g} + \mathbf{e}$, respectively, where the environmental effects follow a bivariate normal distribution with mean vector $\mathbf{0}$ and variance–covariance matrix $\mathbf{E} = \sigma_\mathrm{E}^2 \left[\begin{smallmatrix} 1 & 0 \\ 0 & 1 \end{smallmatrix}\right]$.

Whether or not the mutant can invade is determined by its invasion fitness, i.e. its geometric growth ratio:

$$w\left(\mathbf{g}_\mathrm{m}, \mathbf{g}\right) = \frac{1}{\sum_{j=1}^{N_\mathrm{R}} R_j} \sum_{j=1}^{N_\mathrm{R}} R_j \frac{\overline{\kappa_j}(\mathbf{g}_\mathrm{m})}{\overline{\kappa_j}(\mathbf{g})}\,, \tag{B1}$$

where

$$\overline{\kappa_j}(\mathbf{g}_\mathrm{m}) = \int_{\mathbb{R}^2} \kappa_j\left(\mathbf{g}_\mathrm{m} + \mathbf{e}\right) f(\mathbf{e}) \mathrm{d}\mathbf{e}\,, \tag{B2}$$

is the expected consumption efficiency of the mutant on resource $j$, averaged over environmental effects on the two traits. Here, $f(\mathbf{e})$ is the probability density function of a bivariate normal distribution with mean vector $\mathbf{0}$ and variance–covariance matrix $\mathbf{E}$.

##### B.1 Directional selection and singular trait values

Invasion fitness $w(\mathbf{g}_\mathrm{m}, \mathbf{g})$ can be used to infer the evolutionary dynamics as a sequence of invasions and substitutions by mutations of small effect (e.g. Taylor, 1989; Christiansen, 1991; Eshel et al., 1997; Geritz et al., 1998; Vincent and Brown, 2005; Rousset, 2004; Otto and Day, 2007; Avila and Mullon, 2023). The selection gradient is

$$\mathbf{S}(\mathbf{g}) = \begin{bmatrix} S_1(\mathbf{g}) \\ S_2(\mathbf{g}) \end{bmatrix} = \begin{bmatrix} \left.\dfrac{\partial w(\mathbf{g}_\mathrm{m}, \mathbf{g})}{\partial g_{1,\mathrm{m}}}\right|_{\mathbf{g}_\mathrm{m}=\mathbf{g}} \\ \left.\dfrac{\partial w(\mathbf{g}_\mathrm{m}, \mathbf{g})}{\partial g_{2,\mathrm{m}}}\right|_{\mathbf{g}_\mathrm{m}=\mathbf{g}} \end{bmatrix}. \tag{B3}$$

Each component $S_k(\mathbf{g})$ gives the direction of selection on the genetic basis of trait $k$ when the other is held constant. A positive value indicates selection favouring an increase in the corresponding trait, whereas a negative value indicates selection favouring a decrease. Genetic contributions to trait values $\mathbf{g}^*$ satisfying $\mathbf{S}(\mathbf{g}^*) = \mathbf{0}$ are called singular.

Whether a population evolving through mutations of small effect converges towards a singular trait value is determined locally by the Jacobian of the selection gradient:

$$\mathbf{J}(\mathbf{g}^*) = \begin{bmatrix} \left.\dfrac{\partial S_1(\mathbf{g})}{\partial g_1}\right|_{\mathbf{g}=\mathbf{g}^*} & \left.\dfrac{\partial S_1(\mathbf{g})}{\partial g_2}\right|_{\mathbf{g}=\mathbf{g}^*} \\ \left.\dfrac{\partial S_2(\mathbf{g})}{\partial g_1}\right|_{\mathbf{g}=\mathbf{g}^*} & \left.\dfrac{\partial S_2(\mathbf{g})}{\partial g_2}\right|_{\mathbf{g}=\mathbf{g}^*} \end{bmatrix}. \tag{B4}$$

Assuming isotropic mutational variation (i.e. mutations in the two traits are independent and have equal variances), the singular genetic contribution $\mathbf{g}^*$ is convergence stable if all eigenvalues of $\mathbf{J}(\mathbf{g}^*)$ have negative real parts, in which case nearby populations evolve towards $\mathbf{g}^*$ (Leimar, 2009). Otherwise, $\mathbf{g}^*$ is a repeller and nearby populations evolve away from it.

##### B.2 Quadratic and correlational selection

Once a population has converged to a singular genetic contribution to trait value $\mathbf{g}^*$, the fate of nearby mutants is determined locally by the curvature of invasion fitness. This curvature is summarised by the Hessian of invasion fitness with respect to the mutant genetic values:

$$\mathbf{H}(\mathbf{g}^*) = \begin{bmatrix} \left.\dfrac{\partial^2 w(\mathbf{g}_\mathrm{m}, \mathbf{g})}{\partial g_{1,\mathrm{m}}^2}\right|_{\mathbf{g}_\mathrm{m}=\mathbf{g}=\mathbf{g}^*} & \left.\dfrac{\partial^2 w(\mathbf{g}_\mathrm{m}, \mathbf{g})}{\partial g_{1,\mathrm{m}} \partial g_{2,\mathrm{m}}}\right|_{\mathbf{g}_\mathrm{m}=\mathbf{g}=\mathbf{g}^*} \\ \left.\dfrac{\partial^2 w(\mathbf{g}_\mathrm{m}, \mathbf{g})}{\partial g_{2,\mathrm{m}} \partial g_{1,\mathrm{m}}}\right|_{\mathbf{g}_\mathrm{m}=\mathbf{g}=\mathbf{g}^*} & \left.\dfrac{\partial^2 w(\mathbf{g}_\mathrm{m}, \mathbf{g})}{\partial g_{2,\mathrm{m}}^2}\right|_{\mathbf{g}_\mathrm{m}=\mathbf{g}=\mathbf{g}^*} \end{bmatrix} \tag{B5}$$

The diagonal elements of $\mathbf{H}(\mathbf{g}^*)$ describe quadratic selection on the genetic basis of each trait when the other trait is held constant: a negative value indicates stabilising selection, whereas a positive value indicates disruptive selection. The off-diagonal elements describe correlational selection: a positive value favours mutant deviations in which the two traits change in the same direction, whereas a negative value favours deviations in which they change in opposite directions (Lande and Arnold, 1983; Phillips and Arnold, 1989).

Because $\mathbf{H}(\mathbf{g}^*)$ is symmetric, its eigenvectors define perpendicular directions in trait space and its eigenvalues give the curvature of invasion fitness. The signs of the two eigenvalues define three selection regimes (e.g. Leimar, 2009; Débarre et al., 2014; Mullon et al., 2016; Geritz et al., 2016).

- **(i) Stabilising selection.** If both eigenvalues are negative, invasion fitness has a local maximum at $\mathbf{g}^*$, and selection is stabilising in every direction. The population therefore remains unimodally distributed around $\mathbf{g}^*$ at mutation–selection balance, although correlational selection can shape the covariance and orientation of the trait distribution.

**(ii) One-axis disruptive selection.** If one eigenvalue is positive and the other is negative, invasion fitness has a saddle point. Selection is disruptive along the eigenvector associated with the positive eigenvalue but stabilising along the perpendicular eigenvector. Polymorphism can occur along the former, producing phenotypic variation concentrated around this direction.

**(iii) Two-axis disruptive selection.** If both eigenvalues are positive, invasion fitness has a local minimum, and selection is disruptive along both eigenvectors. Polymorphism can therefore occur along either direction, allowing phenotypic variation to be maintained across both dimensions of trait space.

These conditions are derived for clonal populations. Similar dynamics occur in sexually reproducing diploid populations with additive effects on phenotype, although sexual inheritance and segregation can impose additional conditions on the maintenance of polymorphism (Kisdi and Geritz, 1999).

##### B.3 Analysis

Substituting invasion fitness (eq. B1) into the selection gradient (eq. B3) gives

$$\mathbf{S}(\mathbf{g}) = -\left(\frac{1}{\sigma_{\mathrm{K}}^2+\sigma_{\mathrm{E}}^2}\right)\mathbf{g}. \tag{B6}$$

The unique singular trait value is therefore $\mathbf{g}^* = \mathbf{0}$. Because the Jacobian (eq. B4) is $\mathbf{J}(\mathbf{0}) = -\mathbf{I}/(\sigma_{\mathrm{K}}^2+\sigma_{\mathrm{E}}^2)$, this singular trait value is always convergence stable.

At $\mathbf{g}^* = \mathbf{0}$, the Hessian of invasion fitness (eq. B5) is

$$\mathbf{H}(\mathbf{0}) = \left(\frac{1}{\sigma_{\mathrm{K}}^2+\sigma_{\mathrm{E}}^2}\right)^2 \begin{bmatrix} \sigma_{\mathrm{R}}^2-\sigma_{\mathrm{K}}^2-\sigma_{\mathrm{E}}^2 & \rho\sigma_{\mathrm{R}}^2 \\ \rho\sigma_{\mathrm{R}}^2 & \sigma_{\mathrm{R}}^2-\sigma_{\mathrm{K}}^2-\sigma_{\mathrm{E}}^2 \end{bmatrix}. \tag{B7}$$

The off-diagonal elements show that the strength of correlational selection increases with the correlation between resource properties, $\rho$. When $\rho = 0$, correlational selection is absent and the Hessian is diagonal. When $\rho > 0$, correlational selection favours deviations in which $g_1$ and $g_2$ change in the same direction.

For $\rho > 0$, the eigenvectors of $\mathbf{H}$ define two perpendicular axes: $g_1 = g_2$, along which the traits change in the same direction, and $g_1 = -g_2$, along which they change in opposite directions. Selection is disruptive along $g_1 = g_2$ when

$$\sigma_{\mathrm{K}}^2 + \sigma_{\mathrm{E}}^2 < (1+\rho)\sigma_{\mathrm{R}}^2, \tag{B8a}$$

and along $g_1 = -g_2$ when

$$\sigma_{\mathrm{K}}^2 + \sigma_{\mathrm{E}}^2 < (1-\rho)\sigma_{\mathrm{R}}^2. \tag{B8b}$$

Because $\sigma_{\mathrm{R}}^2 = \delta_{\mathrm{u}}^2$, these conditions determine whether selection is stabilising, disruptive along one axis or disruptive along two axes.

When correlational selection and environmental variance are absent, such that $\rho = 0$ and $\sigma_{\mathrm{E}}^2 = 0$, both conditions reduce to $\sigma_{\mathrm{K}}^2 < \sigma_{\mathrm{R}}^2$. Selection is then disruptive when niche breadth is narrower than the distribution of available resources, consistent with eq. 29 of Slatkin (1979).

Setting $\sigma_{\mathrm{K}}^2 = 1$ and substituting $\sigma_{\mathrm{R}}^2 = \delta_{\mathrm{u}}^2$ into eqs. (B8) gives

$$\rho > -\frac{\delta_{\mathrm{u}}^2 - 1 - \sigma_{\mathrm{E}}^2}{\delta_{\mathrm{u}}^2} \tag{B9a}$$

and

$$\rho < \frac{\delta_{\mathrm{u}}^2 - 1 - \sigma_{\mathrm{E}}^2}{\delta_{\mathrm{u}}^2}, \tag{B9b}$$

respectively. These analytical boundaries distinguish the three selection regimes shown in fig. 1C; see fig. S1 for confirmation from individual-based simulations of large clonally reproducing populations.

#### C Individual-based simulations

We used an individual-based model with discrete, non-overlapping generations to simulate the joint evolution of two quantitative traits, chromosomal inversions, and genetic dominance. Each diploid individual carries two homologous chromosomes containing $L$ loci. At each locus, one allele affects the two quantitative traits. This allele is characterized by its effects on the two traits and by an affinity value determining the weight of its contribution (section 2.1.3). An individual $i \in \{1,\ldots,N\}$ is therefore characterized by $\{a_{1,ik\ell}, a_{2,ik\ell}, b_{ik\ell}\}$ for locus $k \in \{1,\ldots,L\}$ on the maternal ($\ell = 1$) and paternal ($\ell = 2$) chromosome.

The genetic values of the two traits are obtained by combining the allelic effects carried on the two homologous chromosomes according to the model described in the main text (eq. 1). Environmental effects are then added to obtain the phenotypic values. The environmental effect on each trait is sampled from a normal distribution with mean 0 and variance $\sigma_{\mathrm{E}}^2$.

#### C.1 Resource acquisition and fecundity

At the beginning of each generation, the phenotypes of all individuals are calculated from their genetic values and environmental effects (eq. 1).

Each individual's phenotype determines its attack rate on every available resource according to eq. (A5). The total share of resources acquired by each individual is calculated according to eq. (A6) and converted into its fecundity using eq. (A7).

#### C.2 Reproduction

Offspring are produced by a weighted sampling of parents with replacement, with each individual being weighted by its fecundity. Individuals are hermaphroditic, and mothers and fathers are sampled independently. Consequently, selfing is possible, although it is expected to be rare because of the large population size. Each parental pair produces a single offspring, and this process is repeated until the population reaches its constant size $N$. After reproduction, all adults die and are replaced entirely by the newly produced offspring.

Each offspring inherits one chromosome from each parent, including mutation and recombination during the production of gametes.

#### C.3 Mutation

Mutation at genes replaces the existing allele with a newly mutated allele. The new allele has effects on the two traits that are generated by adding a mutational step sampled from a multivariate normal distribution with mean zero and variance–covariance matrix $\mathbf{M}$ (here taken to be diagonal, $\sigma_{\mathrm{M}}^2 \mathbf{I}_2$) onto the effects of the previous allele (following the continuum-of-alleles model). When adding the mutational step to the allelic effects, any absolute values larger than the maximum are capped to the maximal value, ensuring that $-\delta_{\mathrm{u}}/10 \leq a \leq \delta_{\mathrm{u}}/10$.

When dominance evolution is enabled, the affinity parameter is also modified (independently of the mutation at the allelic effect) by adding a mutational step sampled from a normal distribution with mean zero and variance $\sigma_{\mathrm{B}}^2$. When adding the mutational step to the affinity modifier, any negative values are replaced by 0. Mutations occur independently across genes and across offspring.

##### C.4 Recombination

Recombination occurs through crossover events generated along parental chromosomes before gamete formation. Each position has a fixed probability $r$ of being a recombination crossover. In individuals heterozygous for a chromosomal inversion, crossover events occurring within the inverted region are removed, preventing recombination between the standard and inverted chromosome arrangements. In contrast, when both homologous chromosomes carry the same inversion, recombination proceeds normally because the chromosomes are collinear.

##### C.5 Chromosomal inversions

Chromosomal inversions arise through mutation at a specified rate per gamete. Each new inversion is assigned a length sampled from a Poisson distribution with mean $\bar{L}_{\mathrm{I}}$. To ensure a minimum length of two loci, we consider the size of a new inversion as $2+\Delta$, where $\Delta \overset{\text{law}}{\sim} \mathrm{Pois}(\bar{L}_{\mathrm{I}}-2)$ (where $\bar{L}_{\mathrm{I}} > 2$). This new inversion is positioned randomly along the chromosome. If a newly generated inversion overlaps or is immediately adjacent to an existing inversion on the same chromosome, the two are merged into a single larger inverted variant. Inversions do not directly affect fitness but alter patterns of recombination.

##### C.6 Simulation phases

Each simulation consists of a burn-in period and three successive phases.

During an initial burn-in period of $20N$ generations, all individuals have identical reproductive success. This neutral phase allows quantitative genetic variation to accumulate before natural selection is introduced. At the end of this burn-in period, we set $t = 0$.

The **Baseline** phase begins by making reproductive success depend on resource acquisition. During this phase, only mutations affecting quantitative trait values are permitted.

In the **Inversion** phase, chromosomal inversions are allowed to arise while selection and mutation on quantitative traits continue unchanged.

Finally, in the **Dominance** phase, the introduction of new inversions ceases, whereas mutations affecting allelic dominance become possible in addition to mutations that change allelic effects. Existing inversion polymorphisms are maintained, allowing dominance relationships to evolve between polymorphic sites.

##### C.7 Recorded statistics

Summary statistics are recorded at regular intervals throughout the simulations. For every individual, the phenotypic values of both traits and the numbers of heterozygous and homozygous inversions are recorded. At selected generations, additional quantities are calculated, including the additive genetic variance, standing mutational variance, environmental variance, dominance variance, and the phenotypic effects associated with segregating inversions, following section 2.3.1.

For the larger population used to estimate trait heritabilities (of size $10N$ described in section 2.3.1), all individuals are assigned equal fecundity, and no new mutations occur during the production of offspring. Recombination proceeds normally while respecting the recombination landscape imposed by chromosomal inversions.

#### D Characterization of common trait distributions in the presence of inversions

In this Appendix, we consider in more details the six most common trait distributions that arise after inversions emerge and segregate under additive effects, shown in fig. 4. Throughout, we assume random mating and approximate the equilibrium morph frequencies by assuming equal fecundity to all morphs whose trait values correspond to a resource, while other morphs are assumed to have zero fecundity.

**(i) Monomorphic population** The population consists of a single morph located at the origin, $\mathbf{z}_i = (0,0)$, reflecting the symmetry of the resource distribution around the origin.

**(ii) Two inversions, no overshooting** The population is formed by nine morphs, controlled by two inversions segregating independently, each controlling variation in a different trait. Because of the symmetry of the resource distribution around the origin, we consider the frequency of each inversion as 0.5 and no linkage disequilibrium between them. We thus have the four homozygous morphs, $\mathbf{z}_i = (-\delta_\mathrm{u}, \delta_\mathrm{u})$, $\mathbf{z}_i = (\delta_\mathrm{u}, \delta_\mathrm{u})$, $\mathbf{z}_i = (-\delta_\mathrm{u}, -\delta_\mathrm{u})$ and $\mathbf{z}_i = (\delta_\mathrm{u}, -\delta_\mathrm{u})$, each occurs with frequency 0.0625. Four single-heterozygous morphs, $\mathbf{z}_i = (0, \delta_\mathrm{u})$, $\mathbf{z}_i = (-\delta_\mathrm{u}, 0)$, $\mathbf{z}_i = (\delta_\mathrm{u}, 0)$ and $\mathbf{z}_i = (0, -\delta_\mathrm{u})$, each occurs with frequency 0.125. And finally, the double-heterozygous morph, $\mathbf{z}_i = (0,0)$, occurs with frequency 0.25.

**(iii) Single supergene, no overshooting** The population is formed by three morphs, controlled by a single supergene. Here, due to the symmetry of the resource distribution around the origin, the frequency of the supergene is close to 0.5. Therefore, 25% of individuals have the trait $\mathbf{z}_i = (-\delta_\mathrm{u}, -\delta_\mathrm{u})$, 50% of heterozygous individuals have trait value $\mathbf{z}_i = (0, 0)$, and 25% of individuals have $\mathbf{z}_i = (\delta_\mathrm{u}, \delta_\mathrm{u})$.

**(iv) Two inversions, overshooting** The population consists of nine morphs, controlled by two inversions segregating independently, each affecting a different trait. Homozygotes for the derived arrangement overshoot the range of resource properties in the corresponding trait. To compute the equilibrium frequencies of the morphs, we consider one inversion at a time. The frequencies of this focal inversion are $p_\mathrm{aa}$, $p_\mathrm{aA}$ and $p_\mathrm{AA}$, where AA is the overshooting morph, and the morphs have, on average, relative fecundities close to 1, $1+\delta$ and 0 (since the overshooting morph is not adapted to consume any resource). To calculate $\delta$, the augmented fecundity of the adapted heterozygous compared to the adapted homozygous, we consider that each of these morphs is competing for a similar amount of resource (since $R_1 + R_2 = R_4 + R_3$). Therefore, while the fecundity of $aa$ is proportional to $(R_4 + R_3)/p_\mathrm{aa}$, the fecundity of heterozygotes is proportional to $(R_1 + R_2)/p_\mathrm{aA}$. Comparing these fecundities sets $\delta = \frac{p_\mathrm{aa}}{p_\mathrm{aA}}$. Writing the proportion of $a$ and $A$ gametes produced and under random syngamy, we find that the equilibrium frequencies are $p_\mathrm{aa} = p_\mathrm{aA} = 4/9$ and $p_\mathrm{AA} = 1/9$. Assuming no linkage disequilibrium between the inversions, the frequency of each two-locus genotype is simply the product of the corresponding single-locus genotype frequencies.

The four fully adapted morphs, $\mathbf{z}_i = (-\delta_\mathrm{u}, -\delta_\mathrm{u})$, $\mathbf{z}_i = (-\delta_\mathrm{u}, \delta_\mathrm{u})$, $\mathbf{z}_i = (\delta_\mathrm{u}, \delta_\mathrm{u})$, $\mathbf{z}_i = (\delta_\mathrm{u}, -\delta_\mathrm{u})$, are heterozygous or adapted homozygous at both inversions and therefore each occurs with frequency $\left(\frac{4}{9}\right)^2 = \frac{16}{81} \approx 0.2$. The four morphs overshooting in a single trait, $\mathbf{z}_i = (-\delta_\mathrm{u}, 3\delta_\mathrm{u})$, $\mathbf{z}_i = (\delta_\mathrm{u}, 3\delta_\mathrm{u})$, $\mathbf{z}_i = (3\delta_\mathrm{u}, -\delta_\mathrm{u})$, $\mathbf{z}_i = (3\delta_\mathrm{u}, \delta_\mathrm{u})$, each occurs with frequency $\frac{4}{9} \times \frac{1}{9} = \frac{4}{81} \approx 0.05$. Finally, the morph overshooting in both traits, $\mathbf{z}_i = (3\delta_\mathrm{u}, 3\delta_\mathrm{u})$, occurs with frequency $\left(\frac{1}{9}\right)^2 = \frac{1}{81} \approx 0.01$.

**(v) Single supergene, with overshooting** The population is formed by three morphs, controlled by a single supergene. However, the presence of the overshooting supergene breaks the symmetry of the morphs around the origin, and the frequency of the supergene is no longer 50%. To compute the equilibrium frequencies of the morphs $p_\mathrm{aa}$, $p_\mathrm{aA}$ and $p_\mathrm{AA}$, where AA is the overshooting morph, we consider the morphs with relative fecundities 1, $1+\delta$ and 0 (since the overshooting morph is not adapted to consume any resource). To calculate $\delta$, the augmented fecundity of the adapted heterozygous compared to the adapted homozygous, we consider that each of these morphs is competing for a similar amount of resource (recall $R_1 = R_4$). Therefore, while the fecundity of $aa$ is proportional to $R_4/p_{\text{aa}}$, the fecundity of heterozygotes is proportional to $R_1/p_{\text{aA}}$. Comparing these fecundities we find that $\delta = \frac{p_{\text{aa}}}{p_{\text{aA}}}$. Writing the proportion of $a$ and $A$ gametes produced and under random syngamy, we find that the equilibrium frequencies are $p_{\text{aa}} = p_{\text{aA}} = 4/9$ and $p_{\text{AA}} = 1/9$. Therefore, we have about 44.5% of individuals as $\mathbf{z}_i = (-\delta_{\text{u}}, -\delta_{\text{u}})$, 44.5% of heterozygous individuals with trait value $\mathbf{z}_i = (\delta_{\text{u}}, \delta_{\text{u}})$, and 11% of individuals overshoot with a trait value $\mathbf{z}_i = (3\delta_{\text{u}}, 3\delta_{\text{u}})$.

**(vi) Two supergenes, overshooting** The population consists of nine morphs, controlled by two supergenes segregating independently. While one supergene increases both trait values, the other increases one trait and decreases the other trait. This results in a "diamond-like" configuration, and individuals that are homozygotes for both supergenes have trait values that overshoot the resource property range. Therefore, there are four well-adapted morphs that are heterozygous for a supergene and homozygous for the other, $\mathbf{z}_i = (-\delta_{\text{u}}, -\delta_{\text{u}})$, $\mathbf{z}_i = (-\delta_{\text{u}}, \delta_{\text{u}})$, $\mathbf{z}_i = (\delta_{\text{u}}, \delta_{\text{u}})$ and $\mathbf{z}_i = (\delta_{\text{u}}, -\delta_{\text{u}})$, each occurs with frequency 0.125%. Four homozygous morphs that overshoot, $\mathbf{z}_i = (0, 4\delta_{\text{u}})$, $\mathbf{z}_i = (0, -4\delta_{\text{u}})$, $\mathbf{z}_i = (4\delta_{\text{u}}, 0)$ and $\mathbf{z}_i = (-4\delta_{\text{u}}, 0)$, each occurs with frequency 0.0625%. And finally, a single morph that is double heterozygous for the two supergenes, $\mathbf{z}_i = (0, 0)$ is at frequency 0.25.

#### E Metrics of stability for the G-matrix

For each simulation (i.e., parameter combination), we quantified temporal fluctuations in the shape, size, and orientation of the **G**-matrix. These properties were recorded every 100 generations during the last 20′000 generations of each simulation phase (labelled as "Baseline", "Inversions", and "Dominance" in examples in fig. 2).

We then calculated stability metrics within sliding windows of 1′000 generations, each containing 10 observations. Within each window, fluctuations in shape and orientation were quantified as the standard deviations $\text{sd}(\epsilon_{\mathbf{G}})$ and $\text{sd}(\varphi_{\mathbf{G}})$ amongst the 10 observations, respectively. Fluctuations in size were quantified using the coefficient of variation, $\text{cv}(\Sigma_{\mathbf{G}})$, following Jones et al. (2003), since size is an unbounded dimensional metric (i.e., unlike $\epsilon$ and $\varphi$, $\Sigma$ depends on the choice of units of the traits).

**Comparison between presence and absence of inversions** We classified each 1′000-generation window as containing a segregating inversion if at least one inversion segregated at 10–90% frequency throughout the window (i.e. across all 10 generations observed), and as containing no inversion if no inversion segregated during the window. Windows containing segregating inversions for less than 10 generations were removed to exclude transient destabilization (as an illustrative example of this transient effect, see left column of fig. 2 for $\varphi_{\mathbf{G}}$).

For each simulation, we denoted the average of the stability metrics across all windows with inversions by a subscript I, and across all windows without inversions by a subscript N. For example, $\mathrm{sd_I}(\epsilon_{\mathbf{G}})$ and $\mathrm{sd_N}(\epsilon_{\mathbf{G}})$ denote the mean fluctuations in shape within $1'000$-generation windows in the presence and absence of inversions, respectively. The corresponding quantities were also calculated for $\mathrm{cv}(\Sigma_{\mathbf{G}})$ and $\mathrm{sd}(\varphi_{\mathbf{G}})$.

Finally, we computed the relative changes in the temporal fluctuations of the shape parameter in the presence and absence of inversions by

$$\Delta_{\mathrm{I}}\mathrm{sd}(\epsilon_{\mathbf{G}}) = \frac{\mathrm{sd_I}(\epsilon_{\mathbf{G}}) - \mathrm{sd_N}(\epsilon_{\mathbf{G}})}{\mathrm{sd_N}(\epsilon_{\mathbf{G}})}\,. \tag{E1}$$

Analogous relative changes were calculated for matrix size and orientation. Positive values therefore indicate greater temporal fluctuations in the presence of inversions, whereas negative values indicate greater stability.

**Comparison between presence and absence of dominance** First, we discarded all simulations where the average size of the **D**-matrix was less than 10% that of the total genetic variance $\mathbf{G}_{\mathrm{T}}$ during the end of the last simulation phase “Dominance” ($280'000 \leq t \leq 300'000$). For all simulations where dominance evolved, we calculated the stability metrics within the additive phase ($180'000 \leq t \leq 200'000$), during which all effects were constrained to be additive, and the dominance phase ($280'000 \leq t \leq 300'000$), i.e. after dominance had evolved. Metrics were calculated amongst 10 observations within sliding windows of $1'000$ generations as described above.

For each simulation, we averaged the stability metrics across windows in the additive phase (subscript A) and the dominance phase (subscript D). Thus, $\mathrm{sd_A}(\epsilon_{\mathbf{G}})$ and $\mathrm{sd_D}(\epsilon_{\mathbf{G}})$ denote the mean fluctuations in shape at the end of the additive and dominance phases, respectively. The corresponding quantities were also calculated for matrix size and orientation.

Finally, the relative change in the temporal fluctuations of the shape parameter in the presence and absence of dominance is given by

$$\Delta_{\mathrm{D}}\mathrm{sd}(\epsilon_{\mathbf{G}}) = \frac{\mathrm{sd_D}(\epsilon_{\mathbf{G}}) - \mathrm{sd_A}(\epsilon_{\mathbf{G}})}{\mathrm{sd_A}(\epsilon_{\mathbf{G}})}\,. \tag{E2}$$

Analogous relative changes were calculated for matrix size and orientation. Positive values indicate greater temporal fluctuations after the evolution of dominance, whereas negative values indicate greater stability.

The results of these analyses are presented in fig. 6, left column comparing the presence versus absence of inversions, and right column comparing the presence versus absence of dominance.

### Supplementary Figures

for

### The effects of supergene evolution on the structure and stability of the G-matrix

Vitor Sudbrack[2] and Charles Mullon

#### List of Supplementary Figures

Fig. S1 Branching conditions in large haploid population

Fig. S2 Test for multimodality in the absence of inversions

Fig. S3 Effects of inversion evolution on trait heritabilities and genetic correlations

Fig. S4 Phenotype distributions at $t = 200'000$

Fig. S5 Comparing the supergenes in the absence ($t = 200'000$) and presence ($t = 300'000$) of genetic dominance

Fig. S6 Phenotype distributions at $t = 300'000$

Fig. S7 Temporal fluctuations in the shape, size, and orientation of the **G**-matrix as a function of its structure

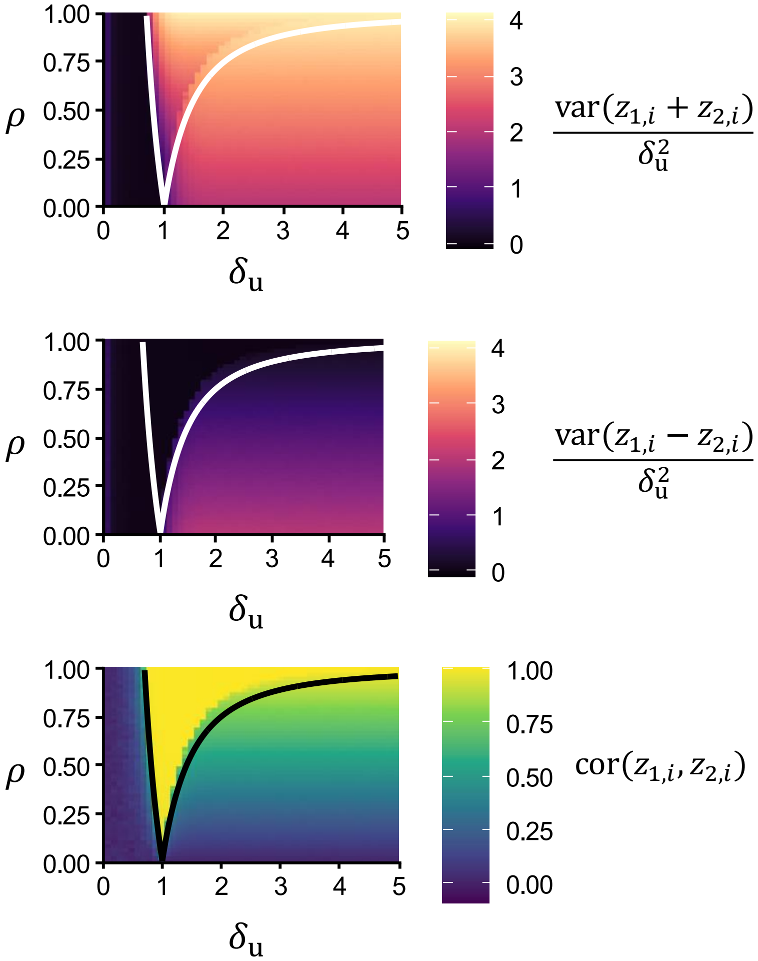


**Supplementary Figure S1: Branching conditions in large haploid population.**
Phenotypic variances along the eigenvectors of the Hessian matrix in a large haploid population: $\text{var}(z_{1,i} + z_{2,i}) = V_{\text{P},1} + V_{\text{P},2} + 2C_{\text{P},12}$ (top) and $\text{var}(z_{1,i} - z_{2,i}) = V_{\text{P},1} + V_{\text{P},2} - 2C_{\text{P},12}$ (middle), as well as the correlation between traits $z_{1,i}$ and $z_{2,i}$, $\text{cor}(z_{1,i}, z_{2,i}) = C_{\text{P},12}(V_{\text{P},1}V_{\text{P},2})^{-1/2}$ (bottom). These variances are normalized with respect to the variance in resource properties, $\sigma_{\text{R}}^2 = \delta_{\text{u}}^2$. Solid lines represent the conditions for the selection regimes described in sec. 2.1.2. Parameters: single haploid locus simulations, $N = 5'000, \sigma_{\text{M}}^2 = 0.001, \mu = 10^{-2}$ and $\sigma_{\text{E}}^2 = 0$, other parameters as default in Table 1.

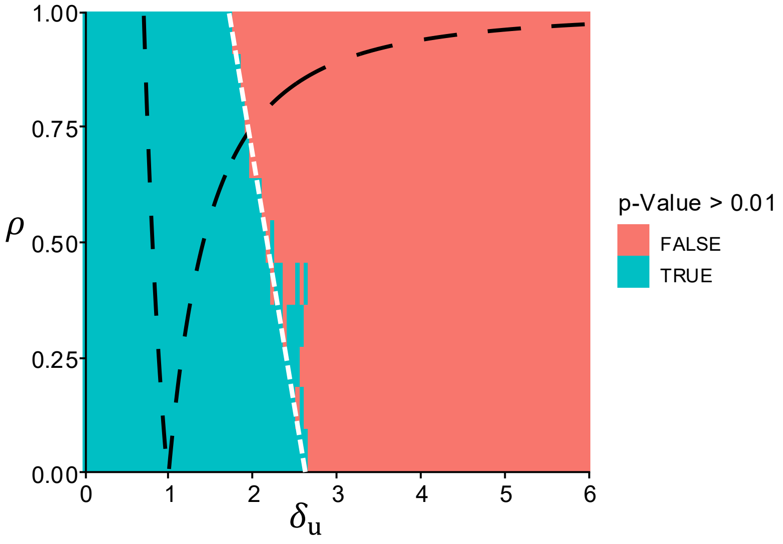


**Supplementary Figure S2: Test for multimodality in the absence of inversions.**
The $p$-value from Hartigan's dip test, a nonparametric test for unimodality that measures the maximum difference between the observed distribution and the best-fitting unimodal distribution, was used to assess whether the phenotype distribution in the absence of inversions ($t = 100'000$) deviated from unimodality. Distributions with $p < 0.01$ (red) were classified as significantly multimodal, whereas those with $p \geq 0.01$ (blue) were considered consistent with unimodality. Multimodal phenotype distributions in the absence of inversions indicate that selection was strong enough to generate linkage disequilibrium between alleles with similar phenotypic effect. We have neglected these simulations during our study by adding a white ribbon to figures (the left-most boundary of the ribbon is represented by the white dashed line), since in these scenarios inversions are not needed to generate physical linkage and niche diversification in the first place. Fixed parameters: same as fig. 2.

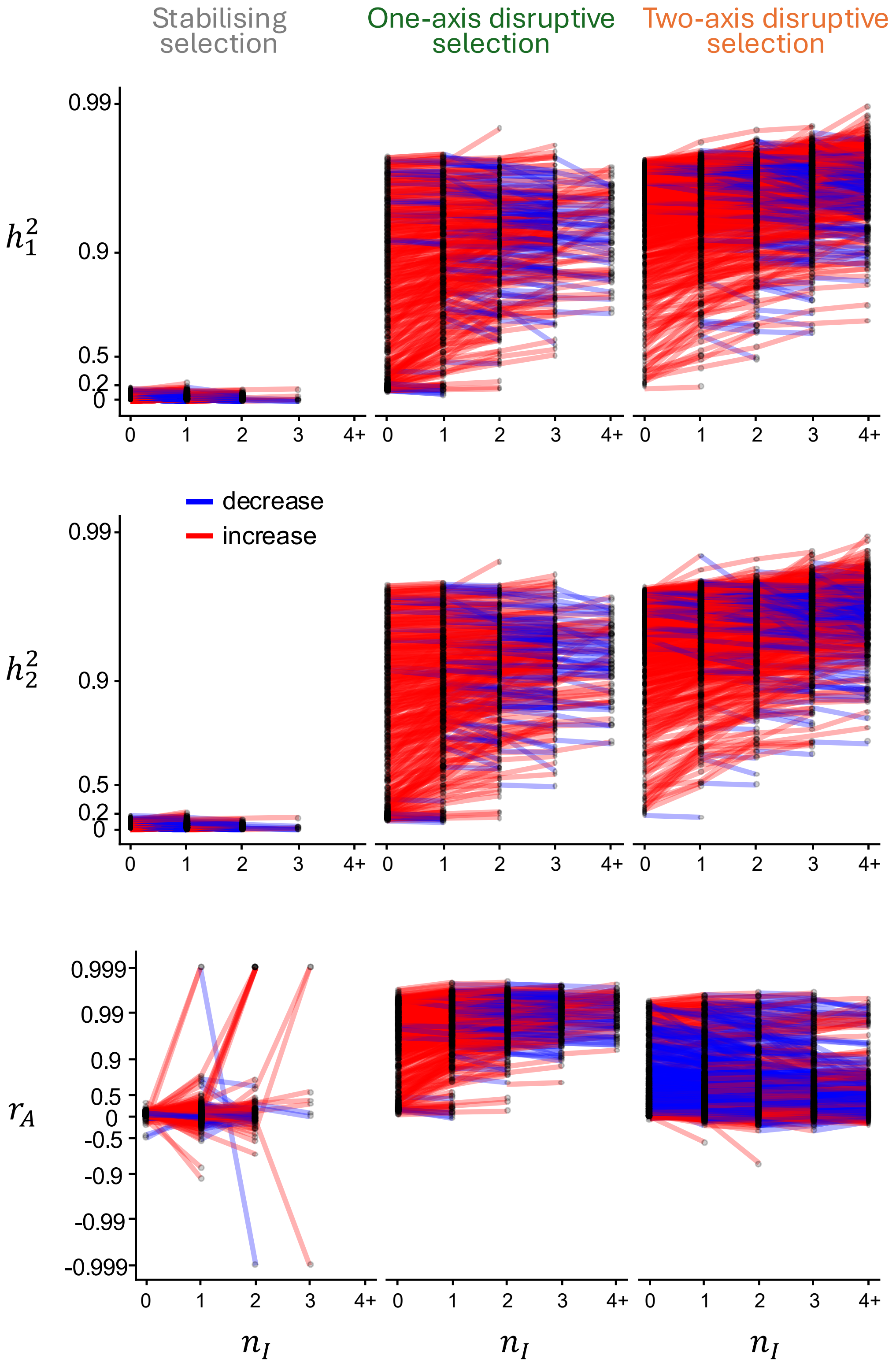


**Supplementary Figure S3: Effects of inversion evolution on trait heritabilities and genetic correlations.** Heritability of trait 1 ($h_1^2$; top), heritability of trait 2 ($h_2^2$; middle), and the additive genetic correlation between traits ($r_A$; bottom) as a function of the number of inverted strata ($n_I$) segregating at intermediate frequencies ($0.1 < p < 0.9$) up to generation $t = 200'000$. Each line connects successive values of the same simulation before and after the addition of an inverted stratum. Segments are coloured red when the addition of a stratum increases the corresponding heritability (or $|r_A|$) and blue when it decreases it. Simulations are grouped by selection regime according to the analytical conditions derived in sec. 2.1.2: stabilising selection (left), one-axis disruptive selection (centre), and two-axis disruptive selection (right). Fixed parameters are as in fig. 2.

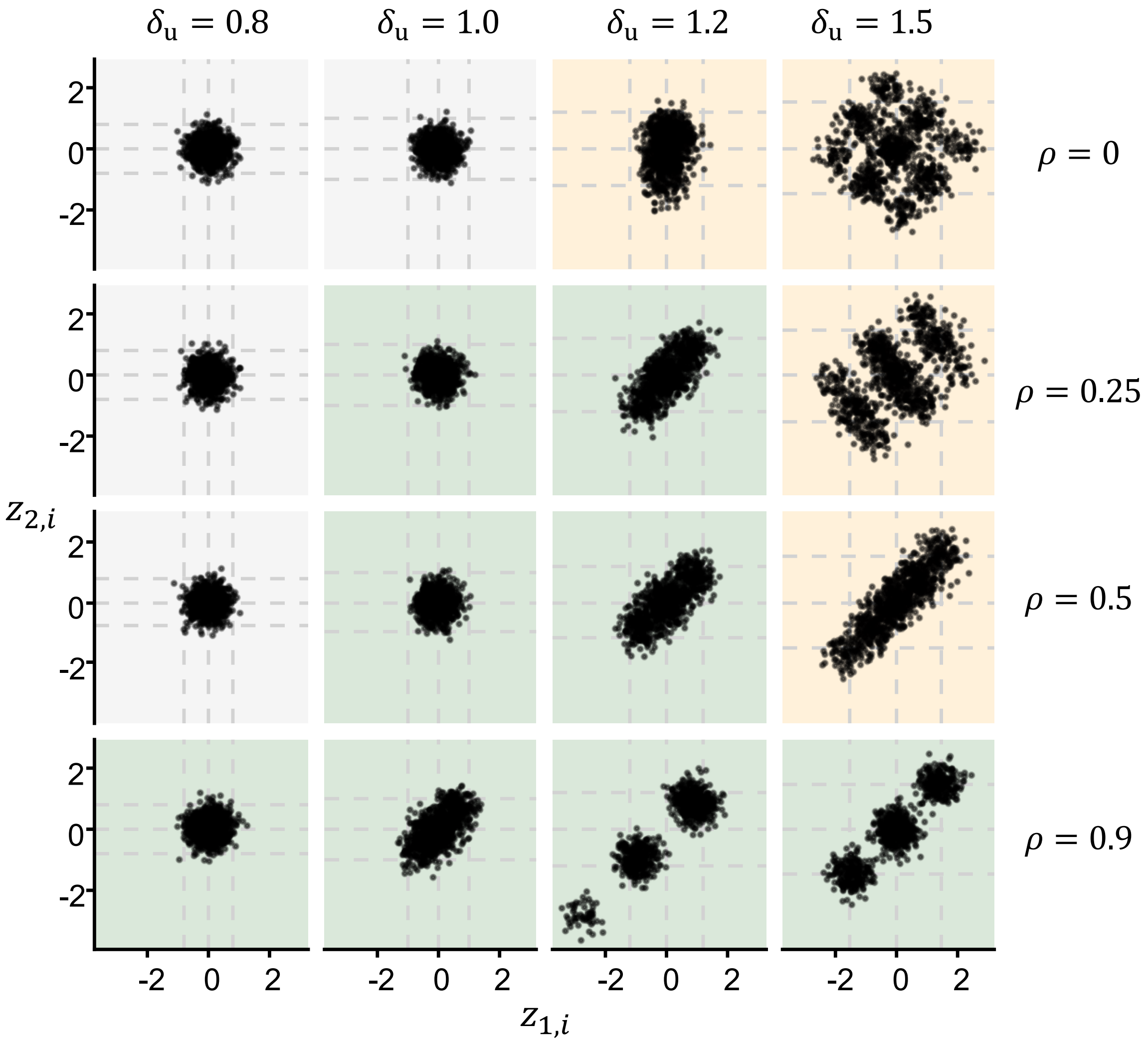


**Supplementary Figure S4: Phenotype distributions at** $t = 200'000$**.**
Phenotypic values for individuals in the population for different values of the resource divergence, $\delta_u$, and resource correlation, $\rho$ (columns and rows, respectively) at $t = 200'000$ (end of phase B in fig. 2). The background colour indicates the predicted selection regime: stabilising selection (gray), one-axis disruptive selection (green), and two-axis disruptive selection (orange). All other parameter take the default value in Table 1.

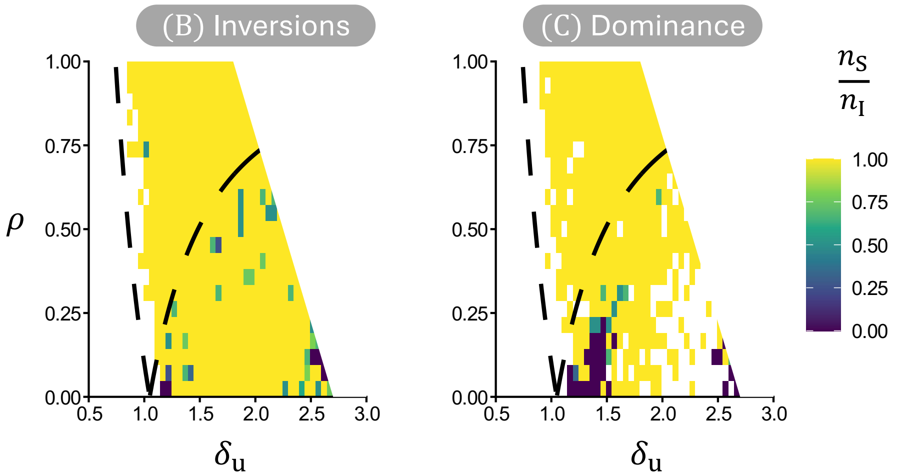


**Supplementary Figure S5: Comparing the supergenes in the absence ($t = 200'000$) and presence ($t = 300'000$) of genetic dominance.**
Fraction of inverted strata that are supergene (i.e., with phenotypic effects in both trait values), $n_S/n_I$ when all phenotypic effects are additive ($t = 200'000$, left) and after the evolution of genetic dominance ($t = 300'000$, right panel). Bright colours indicate the presence of supergenes. In the right panel, only simulations where the size of the **D**-matrix was at least 10% of the total genetic variance $\mathbf{G}_T$ are present. Fixed parameters: same as fig. 2.

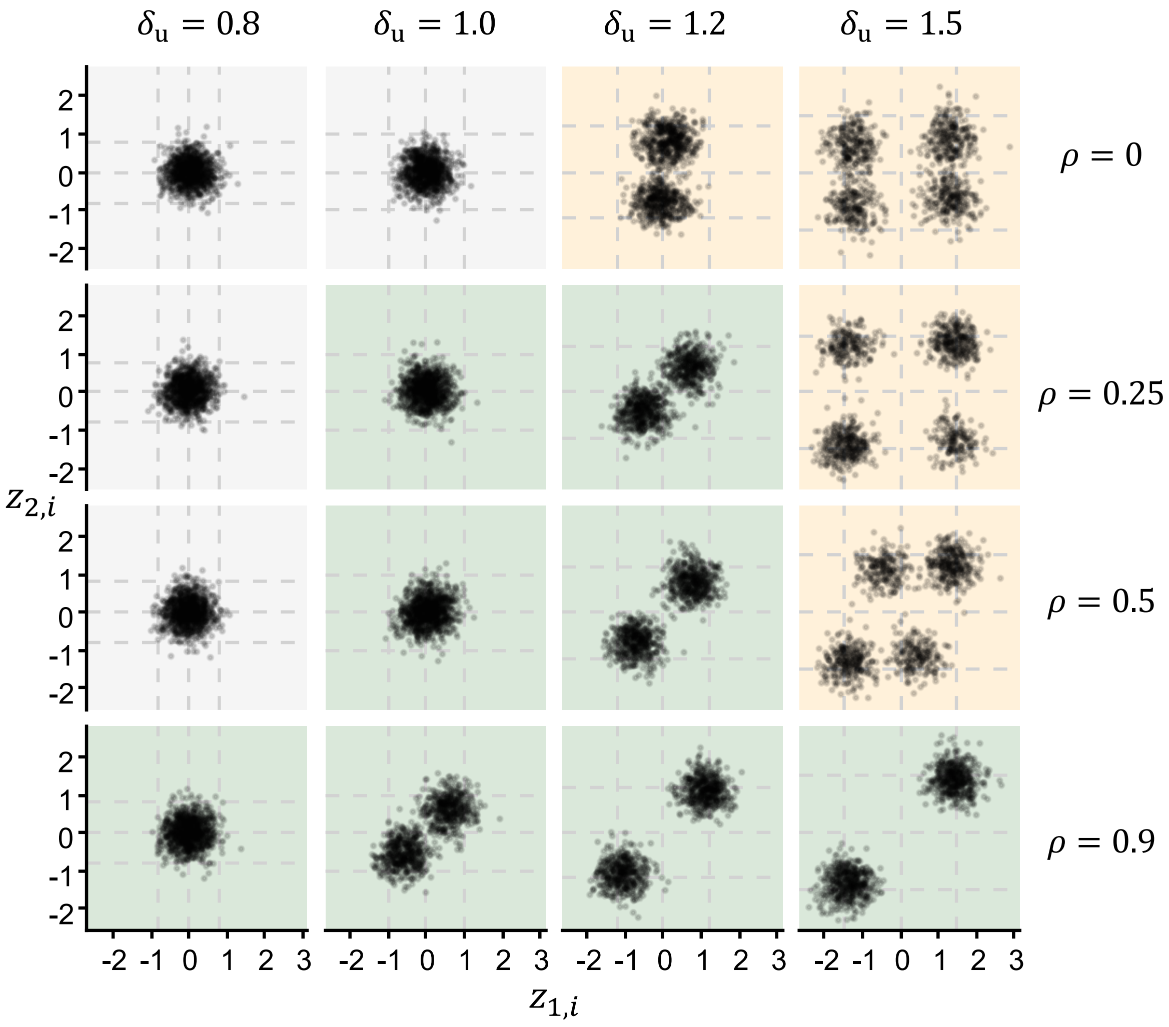


**Supplementary Figure S6: Phenotype distributions at** $t = 300'000$**.**
Phenotypic values for individuals in the population for different values of the resource divergence, $\delta_u$, and resource correlation, $\rho$ (columns and rows, respectively) at $t = 300'000$ (end of phase C in fig. 2, after the evolution of genetic dominance). The background colour indicates the predicted selection regime: stabilising selection (gray), one-axis disruptive selection (green), and two-axis disruptive selection (orange). All other parameter take the default value in Table 1.

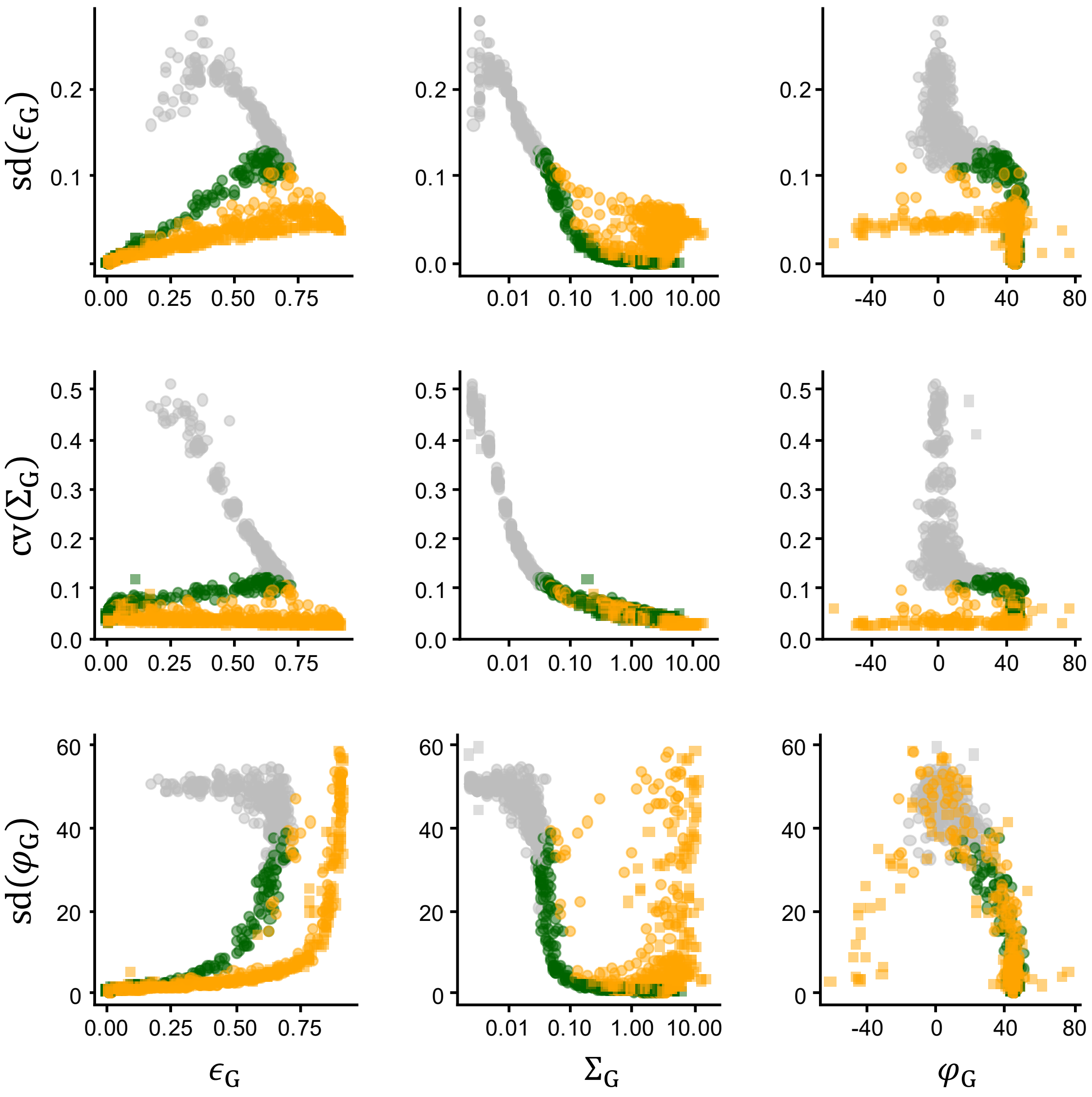


**Supplementary Figure S7: Temporal fluctuations in the shape, size, and orientation of the G-matrix as a function of its structure.**

Temporal fluctuations in the shape (top row), size (middle row), and orientation (bottom row) of the **G**-matrix as a function of its average shape ($\epsilon_{\mathbf{G}}$; left column), size ($\Sigma_{\mathbf{G}}$; middle column), and orientation ($\varphi_{\mathbf{G}}$; right column). Fluctuations and averages were quantified within 1′000-generation windows (see Appendix E for details). Circles indicate windows without segregating inversions, whereas squares indicate windows containing at least one segregating inversion. Colours indicate the selection regime: stabilising selection (gray), disruptive selection along the correlated axis (green), and disruptive selection along both axes (orange). Fixed parameters: same as fig. 2.
